# Halting predicted vertebrate declines requires tackling multiple drivers of biodiversity loss

**DOI:** 10.1101/2025.01.02.630023

**Authors:** Pol Capdevila, Duncan O’Brien, Valentina Marconi, Thomas F. Johnson, Robin Freeman, Louise McRae, Christopher F. Clements

**Author notes:** These authors contributed equally to this work (Pol Capdevila, Duncan O’Brien, Louise McRae, Christopher F. Clements). Previous address (Pol Capdevila).

## Abstract

Anthropogenic threats are reshaping Earth’s biodiversity at an unprecedented rate and scale^1–3^. Conservation policies often prioritise threats like habitat loss and exploitation based on their global prevalence. However, these assessments rarely quantify the impacts of individual or interacting threats, potential masking the true effects of the Anthropocene^4–6^. Here, we quantitatively analyse the trends of 3,129 vertebrate populations worldwide with documented exposure to specific and multiple threats. Populations impacted solely by habitat loss or exploitation, the most prevalent threats, do not show the fastest declines. Rather, populations exposed to disease, invasive species, pollution, and climate change decline more rapidly. However, habitat loss and exploitation – along with climate change – do act as additive interactive threats, amplifying population declines. Notably, these interactive threats contribute to population declines, more than temporal or spatial sources of variation. Finally, counterfactual scenarios show that to achieve global non-negative vertebrate population trends, we need to mitigate the effects of multiple threats. These findings underscore the urgency of addressing the compounding effects of multiple threats to halt biodiversity loss and suggest that the local-scale impacts of climate change may be more severe than previously recognized.

## Introduction

Anthropogenic effects on Earth’s natural ecosystems are now more pervasive than ever^1–3^, with threats such as habitat loss, climatic change, invasive species, pollution, and exploitation jeopardizing more than 1 million species with extinction^3,7,8^. Many international conservation objectives, including the Kunming-Montreal Global Biodiversity Framework^9^ and the Sustainable Development Goals^10^, have been set to halt the loss of biodiversity^11,12^ and protect the services it provides to humanity^13,14^. To achieve these targets, conservation policies often rely on global rankings of threats, which order them according to their reported prevalence, to prioritise management decisions. Conservation reports have identified habitat loss and exploitation as the leading causes of biodiversity loss^1–3^, whilst other threats, such as climate change, are predicted to become increasingly important in the coming decades^2,15^. However, these threat rankings are usually ordered according to global species assessments (e.g. IUCN Red List^1,2^) or pressure maps^4,5^ based on the presence/absence of threats rather than their actual impact on ecological systems (but see^16^). Moreover, global assessments group threat data at the species level, which might not necessarily reflect the status of populations at local scales^17,18^. Consequently, global biodiversity assessments based on these approaches may underestimate the real magnitude of the impacts of global change^19^.

Explicitly accounting for the effects of threats on biodiversity is further complicated by the fact that most species globally (about 80%^2^) are exposed to more than one threat simultaneously^8^. Multiple threats can interact in complex and non-linear ways^20–22^, and their cumulative effects can be higher (synergistic effects) or lower (antagonistic effects) than the sum of their individual effects (additive effects)^20,21^. Synergistic interactions are of particular concern for ecology and conservation^21,23^, because they have the potential to further accelerate biodiversity loss by amplifying the impacts of multiple threats^22,24^. Identifying the prevalence of synergic, antagonistic, or additive interactions is therefore critical for prioritizing conservation actions^23^, as well as gauging the true impacts of global change on biodiversity. While some studies emphasize the prevalence of synergies as a major cause of biodiversity loss^22^, others suggest that additive effects are more common^21,23^. However, the difficulties of accounting for multiple stressors in the analysis of real-world data mean that such studies are largely based on meta-analyses of experimental studies^21,23^, with limited temporal and spatial scales. These constraints might render an incomplete picture of the impacts of multiple threats and precludes identifying which threats are more likely to cause synergistic interactions. In fact, conservations efforts are known to improve biodiversity but often are insufficient to halt biodiversity decline^25^ possibly due to many efforts targeting singular threats. To date, no study has used empirical data to examine the prevalence of the different types of threat interactions at a global scale and their association with population trends^23^.

Here we show how information on threats to wildlife populations can both provide a more informative estimate of biodiversity trends and guide approaches to reverse the declines. We quantify the trends of populations exposed to single and multiple interacting threats to vertebrate populations at a global scale using a subset (3,129) of time-series from the Living Planet Database^26^ (LPD hereafter; Figure. 1). This dataset contains information about which threats are impacting each population according to the data sources (e.g., scientific manuscripts, technical reports). Threats were classified into five broad categories – exploitation, habitat loss, disease, pollution, climate change, or invasive species, matching those of the IUCN threat categories^8^ (for comparability with previous studies), as well as a category of “no threat” exposure. To determine which threats, and threat combinations are associated with the most dramatic declines we estimated the population trends exposed to single and multiple threats using multilevel Bayesian models, accounting for spatial and temporal autocorrelation^27^. To determine the prevalence of synergistic, antagonistic, and additive interactions, we compared the estimated population trends for interactive threats versus their additive combinations (Figure S1). In brief, our model structure allows us to build threat combinations (i.e. absent, additive or interactive) in an incremental fashion, with an additive threat combination contributing to the interactive estimate (see Methods). Finally, to assess the potential benefits of management actions, we developed counterfactual scenarios describing how the removal of single or multiple threats would alter the predicted global trends of vertebrate populations.

**Fig. 1.**
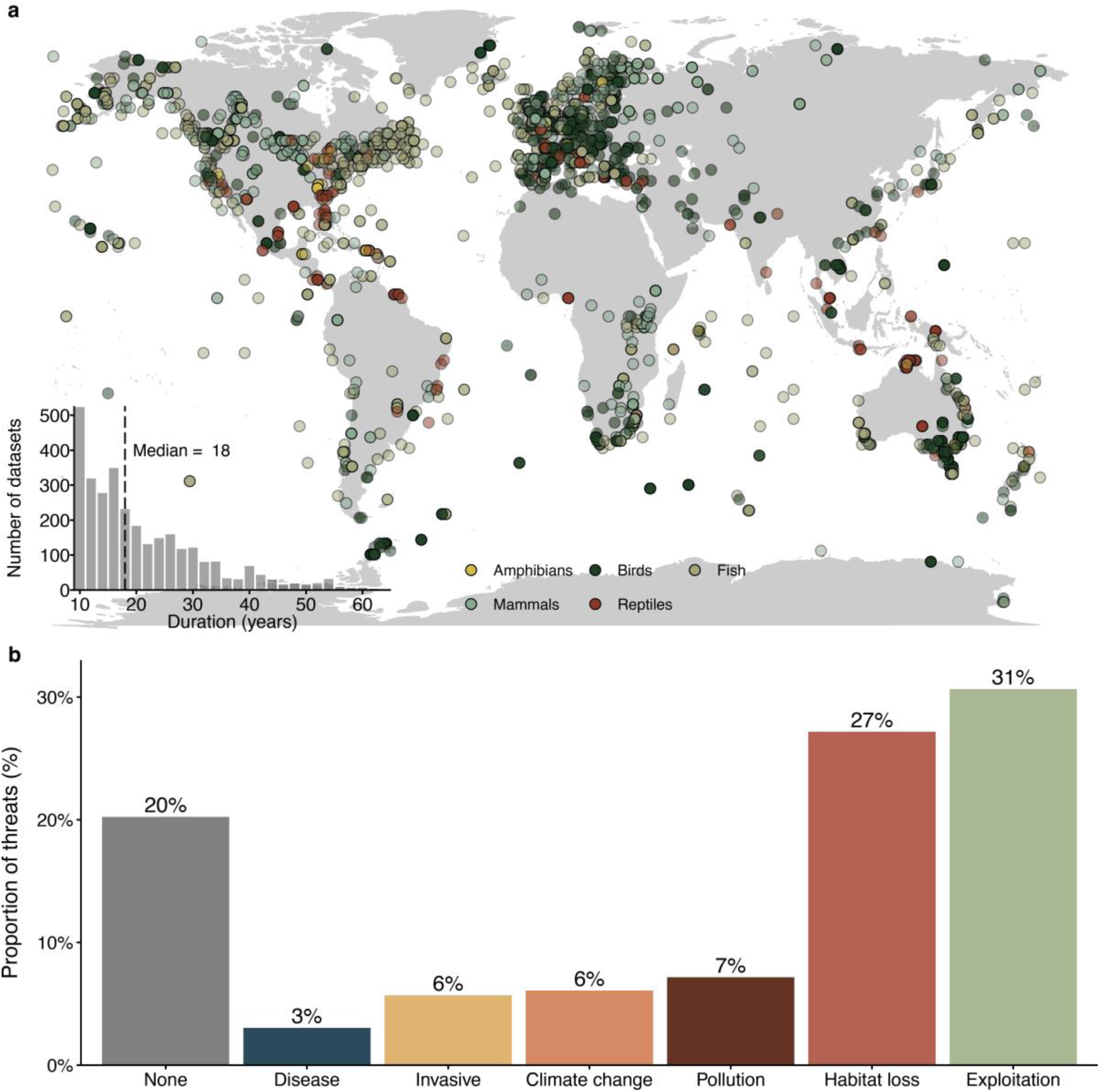
Habitat loss and exploitation are the most prevalent threats impacting vertebrate populations. (**a**) Spatial distribution of the population trends from 3,129 population time-series from 1,281 species contained in the Living Planet Database. (**b**) Proportion of populations affected by the different threats. 20.00% of the populations are not exposed to threats, 3.04% are exposed to disease, 5.69% to invasive species, 6.07% to climate change, 7.16% to pollution, 27.20% to habitat loss and 30.60% to exploitation.

## Habitat loss and exploitation are not associated with the fastest population declines

Although habitat loss and exploitation are the most prevalent threats^1,2^, and are often the main targets of conservation^10,28^, our results show that they are not associated with the fastest population declines. Our multilevel Bayesian models show that being exposed to any threat accelerates the decline of vertebrate populations worldwide (Figure 2**a**; Table S1). For instance, vertebrate populations exposed to invasive species decline five times faster (-6.01% yr^-1^, 95% credible interval (CI) = [-30.76% yr^-1^, 10.10% yr^-1^]) than those not exposed to any threat (1.70% yr^-1^, CI = [-13.64% yr^-1^, 20.00% yr^-1^]; Figure 2**a**). Similarly, despite being much less prevalent threats (Figure. 1**b**), pollution (-1.79%yr^-1^ [-17.51-17.05%]), climate change (-3.14% yr^-1^ CI=[-17.21%, 17.05%]), and disease (-4.63 yr^-1^ CI=[-14.89%, 6.49%]) also associate with population declines (Figure 2**a**). Finally, contrary to our initial expectations, exploitation (-0.85%yr-1 CI=[-15.47, 15.43]) and habitat loss (-0.42%yr-1 CI=[-17.04, 18.71]) are the threats associated with the slowest population declines (Figure 2**a**).

**Fig. 2.**
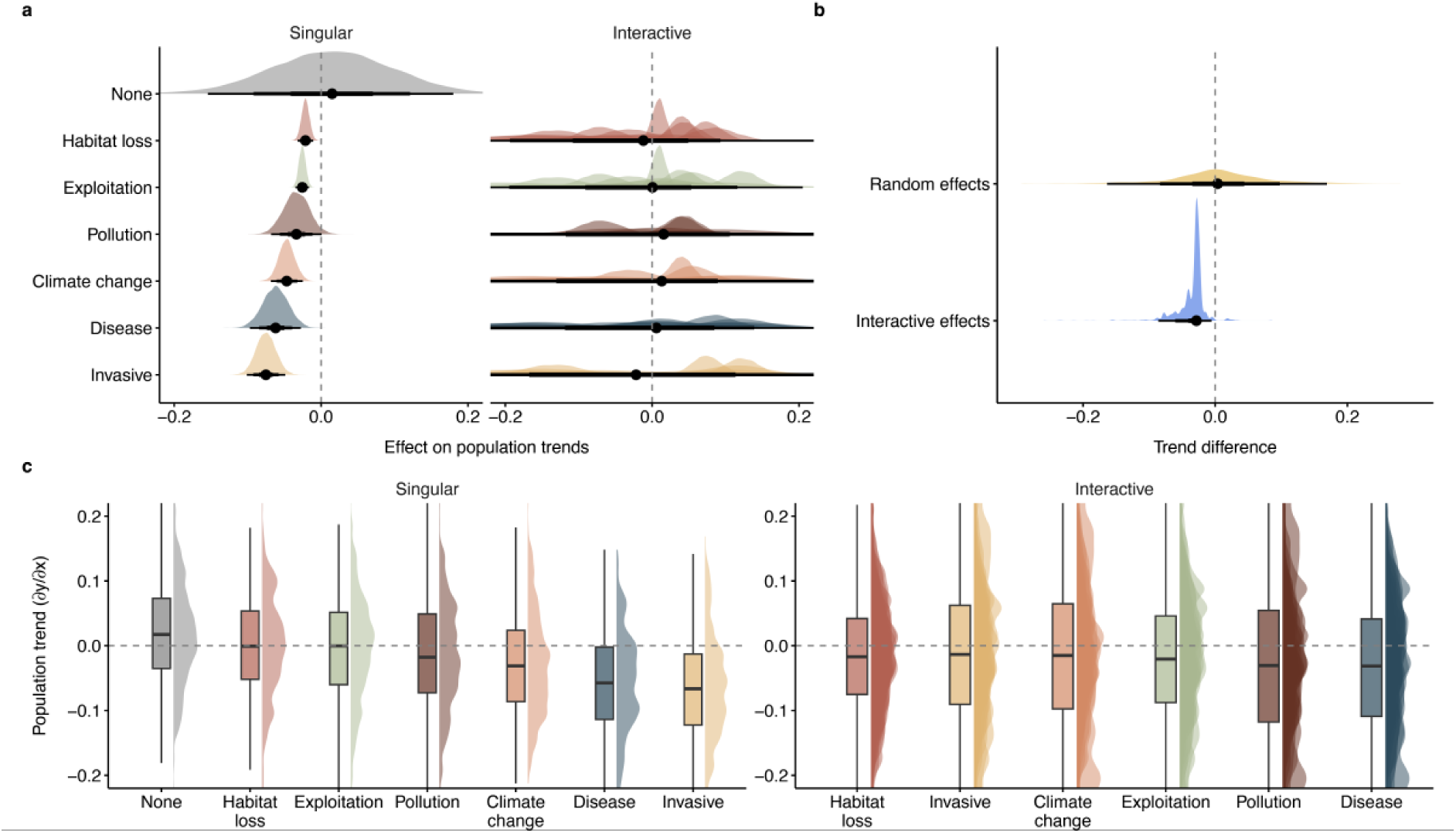
Invasive species, disease and climate change accelerate the decline of vertebrate populations. (**a**) Effect of threats on vertebrate population trends of 3,129 vertebrate population time-series from 1,281 species contained in the Living Planet Database. “None” represents the estimated trend in the absence of threats. Less prevalent threats such as invasive species, disease, or climate change, have a larger effect on vertebrate population trends than more pervasive threats such as habitat loss or exploitation. Singular threats indicate the effect of the threats when acting in isolation. Interactive threats indicate those acting in conjunction with other (one or two) threats. (**b**) Distribution of the influence of the random effects and the presence of threat interactions on the population trends. To estimate the difference in population trends, we first estimated posterior draws from the fitted model without considering the threat interactions, nor the random effects (base predictions). We repeated these predictions with including the random effects and the interactive threats, separately. Then subtracted the two forementioned posterior draws datasets from the population trends of the base predictions. (**c**) Distribution of estimated trends given the realisation of the combined model coefficients presented in (**a**). Trends are calculated as the first derivative of the estimated model slope(s). All density plots are based on 1000 samples from the posterior distribution of the coefficients (**a**) or trend estimates (**b,c**). The reported values are the highest posterior density median values (circles), with 50% (thickest bars), 80% and 95% (thinnest bars) uncertainty intervals.

The interactive threats displayed a much wider range of trajectories than single ones, showing the most positive and negative trends (Figure 2**a**). Pollution (-2.83%yr-1 CI=[-44.96, 50.77]) and climate change (-1.42%yr-1 CI=[--30.06, 33.70]) are the interactive threats displaying the largest variability (Figure 2**a**), while the population trends displaying the fastest declines (Table S1) are those experiencing the combined influence of pollution, climate change and disease (-16.74% yr^-1^ CI=[-31.55, 1.00]), and the interaction between pollution and invasive species (-10.27% yr^-1^ CI=[-67.11, 134.32]). However, the latter is extremely uncertain given its limited representation in our dataset (Figure 1). In terms of average trends, populations experiencing pollution (-2.83%yr^-1^ CI=[-44.96, 50.77]) and disease (-2.97%yr^-1^ CI=[-23.61, 18.56]) interactions with any other threats show the fastest average decline (Figure 2**a,c**). Overall, these results underscore the complexity of global conservation, emphasising the context-dependent nature of these threats and how their interactions add to the uncertainty of biodiversity trends.

Although vertebrate population trends exhibit considerable uncertainty when assessing the effects of interactive threats (particularly between taxa and systems: Figures S2-S5, Tables S2-S3), some studies suggest this uncertainty may arise from correlative factors, such as spatial and temporal autocorrelation^27^. Therefore, we compared the impact of spatial and temporal autocorrelation (random effects) with the overall effect of interactive threats on population trends by estimating their differences. Specifically, we contrasted trend predictions without random effects or threat interactions with those including these factors, and then subtracted the latter from the former (see Figure 2**b** and Methods for further details). Our findings indicate that, despite the variability in population trends, interactive threats are associated with stronger declines than random effects (Figure 2**b**). Therefore, these results provide evidence that interactive threats are contributing to estimated biodiversity loss, more than temporal or spatial sources of variation.

## Synergies are the least frequent interaction type

Contrary to the common perception that synergies are a frequent consequence of global change^22^, our findings suggests that most threats instead interact additively (61%-100%; Figure 3). Here, we tested the prevalence of the additive, synergistic and antagonistic interactions across the global model, as well as system- and taxon- specific models (see Methods). Across the different models and threats most interactive effects are classified as additive (Figure 3 and Figures S6-S8; Tables S4-S6). For instance, when using the global model, additive threats represent 84.90% of interactions involving climate change, 80.60% for disease, 89.20% for exploitation, 85.83% for habitat loss, 78.47% for invasive species, and 85.29% for pollution (Figure 3; Table S4).

**Fig. 3.**
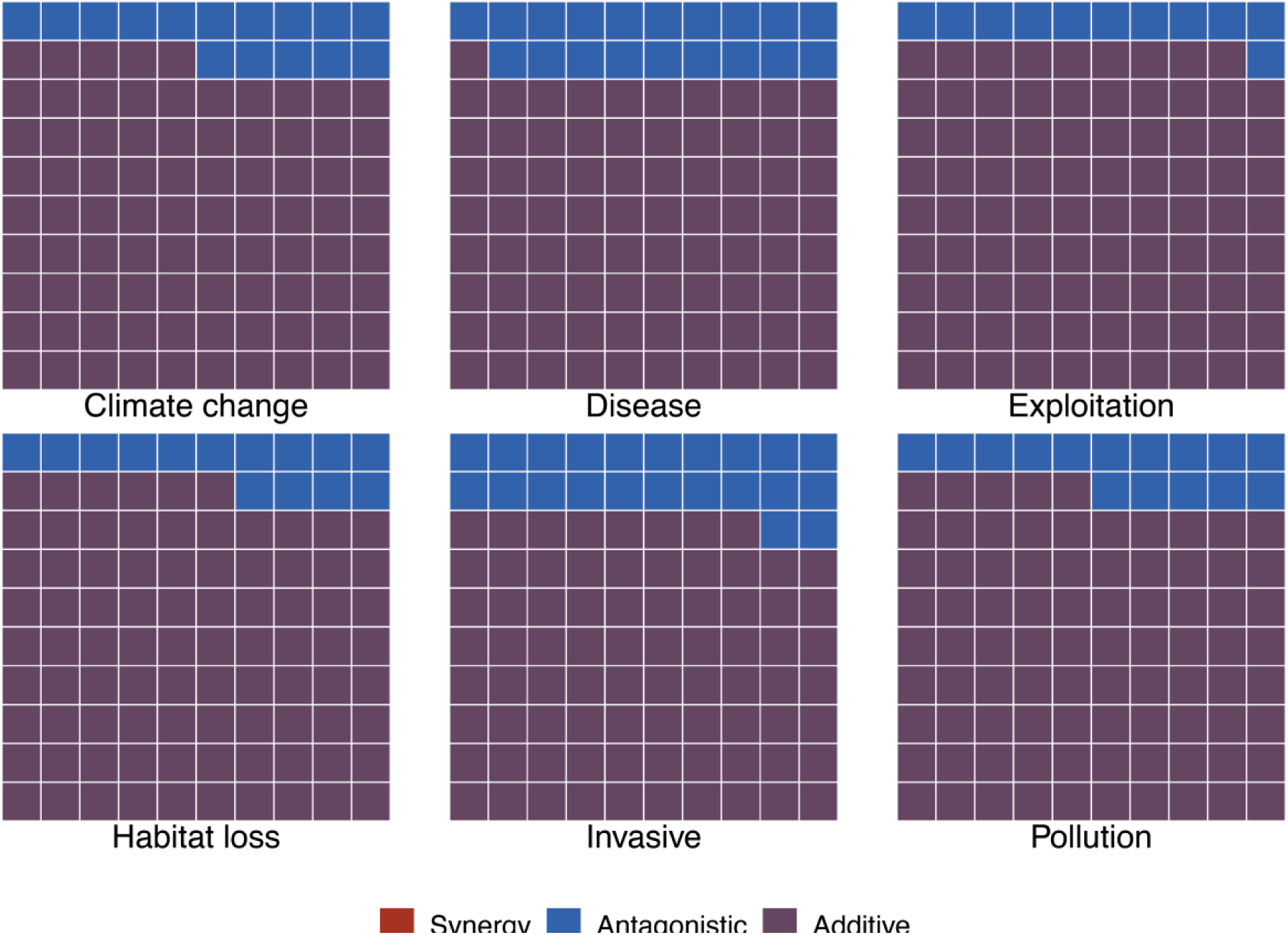
Synergies are not the most common interactive effect across threats. Interactive effects of the different threats. Most interactive effects are additive (purple) or antagonistic. The interactive effects were classified comparing the estimated population trends for interactive threats versus their additive combinations (additions of the population trends exposed to single threats). They were defined as additive, when the trends for populations exposed to the combined threats were equal to the summation of the trends for populations exposed to the individual threats. Synergistic, when the trends for populations exposed to the combined threats were larger than the summation of the trends for populations exposed to the individual threats. Antagonistic, when the trends for populations exposed to the combined threats were smaller than the summation of the trends for populations exposed to the individual threats. No interaction was synergistic in the global model, while 84.90% was additive for climate change, 80.60% for disease, 89.20% for exploitation, 85.80% for habitat loss, 78.40% for invasive species, and 85.30% for pollution.

All non-additive effects in the global model are antagonistic, accounting for a proportion varying between 10% (exploitation) and 22% (invasive species) of the interactions (Figure 3 and S6; Table S4). Synergies do not appear in this global model though the amphibian specific model does display synergistic threats (Figure S7). These findings therefore suggest that, at the macroecological level, synergies are not as prevalent as perceived in the literature^23^.

The prevalence of antagonistic interactions over synergies could be due to multiple reasons (reviewed in^23^). For example, a single threat may decrease the sensitivity of a species to a second threat (e.g. a threat decreasing the size of a population might make easier to avoid the predation from invasive species^29^). However, whilst a high proportion of antagonistic effects could be perceived as a positive outcome, it demonstrates the difficulties of predicting population responses to multiple threats^20–23^.

Still, it is worth highlighting that system and taxa are idiosyncratic in their threat interactions. For example, with amphibians being the only taxonomic group exhibiting synergies as the main non-additive interactions (Figure S7), this is in line with their increasingly threatened status^1,3,30^. Indeed, disease, exploitation and invasive species are the threats associated with the most synergies in amphibians (Figure S7), reflecting their vulnerability to these threats^30^. Terrestrial systems are also the only system to exhibit synergies (Figure S8) though this is likely the contribution of terrestrial amphibians. Despite these results, our estimated dominance of non-additive effects are, most likely, conservative, given the uncertainty in the population trend estimates (see Methods and Figure S6).

## Only mitigating multiple threats will reverse predicted vertebrate population declines

Although exploitation and habitat loss are not the threats associated with the fastest population declines, our counterfactual scenarios show that management actions aimed at tackling them may be more effective at slowing the predicted rate of decline in vertebrate populations than targeting other individual threats (Figure 4 and S9). However, addressing any one threat individually is insufficient to reverse vertebrate population declines, whereas the removal of all threats does, accounting for a 226.44% proportional increase in predicted population trends. This is a key indication that single target interventions are insufficient for reversing global biodiversity declines.

**Fig. 4.**
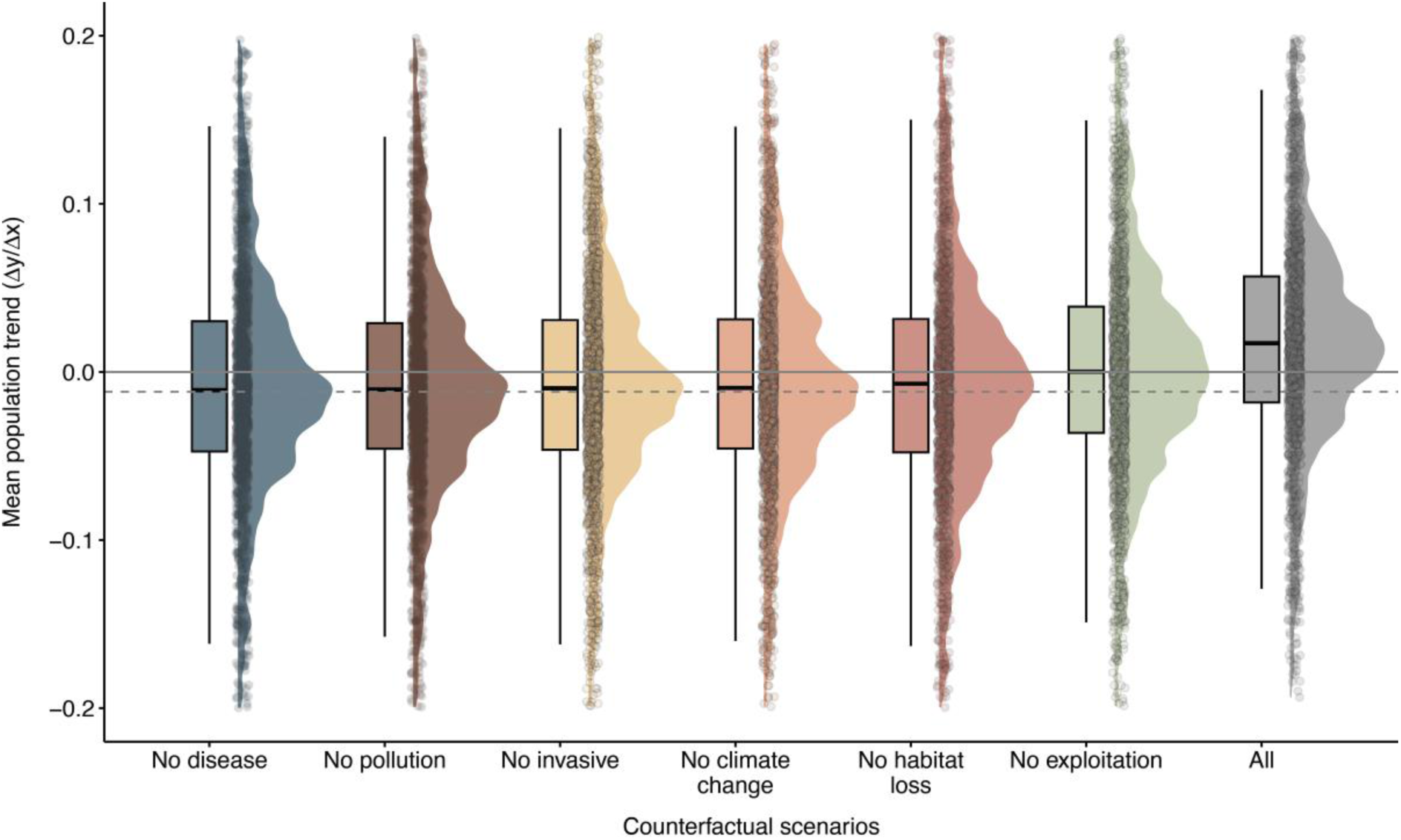
The counterfactual scenarios suggest that lowering exploitation would be the most effective action to slow down vertebrate populations decline. The counterfactual scenarios represent the changes in the population trends of the 1,740 vertebrate time-series affected by threats, had there been no disease, no pollution, no invasive species, no climate change, no habitat loss, no exploitation or no threats. The continuous, grey line represents when the population trend is 0. The dashed grey line is the median trend of the threatened populations without any of the counterfactual scenarios. Box plots depict the distribution of population trends where: the thick line in the middle represents the median value; the lower and the upper bounds of the box represent the 25th–75th quartiles, respectively; and the straight lines represent the minimum to the lower quartile (the start of the box) and then from the upper quartile (the end of the box) to the maximum.

Specifically, these counterfactual scenarios adjust for both the frequency and the potential impact of the threats on vertebrate populations by simulating scenarios where the effects of threats are removed for each of the populations affected by single or multiple threats (see Methods). Removing the effects of exploitation results in the largest increase in predicted population trends (81.62% relative increase), although the median trend is still negative, followed by habitat loss (16.87%; Figure 4). The high prevalence of exploitation and habitat loss makes them more likely to interact with other threats. Consequently, despite their lower impact as single threats, mitigating exploitation and habitat loss should remain a conservation priority at a global scale. In contrast, when the invasive species, pollution and disease threats are removed, there is a smaller effect on the overall global population trends (14.19%, 10.49%, and 6.94%, respectively; Figure 4), either because of their low prevalence or low impact. In addition, for the first time, we show that managing climate change could substantially improve predicted vertebrate populations, highlighting it as a key component of the current biodiversity crisis^15,31^. These findings contrast with previous studies suggesting that the effects of climate change are still modest^1,2,32^, highlighting the importance that climate policies could have in reverting biodiversity loss^33^.

The discrepancy between our results and previous assessments is due to the different methodologies used to rank global threats and highlights the risk of such approaches to prioritise conservation actions^32^. Most global threat analyses are based on information from IUCN Red List assessments (e.g.^1,2,17^), and are estimated using the presence/absence of threats to species rather than the magnitude of their impacts^8^. Therefore, it is not surprising that global analyses often find habitat loss and exploitation as the most prevalent threats, given the millennia history of human’s exploitation and modification of natural ecosystems^34,35^. On the contrary, here, we incorporate both the potential impacts (population trends) and the frequency (their prevalence in our dataset) of threats on natural systems. The IUCN Red List has recently incorporated threat intensity in their criteria^8,16^, which will be key to characterise the impacts of threats and guide conservation priorities.

It is worth noting, however, that population trend estimates show large uncertainties, so our findings should be interpreted with caution. In addition, the counterfactual scenarios presented here assumed that the cause of the population trends are threats. However, we are interpreting the counterfactual scenarios as the “influence of threats on predicted trends” only – i.e they represent tentative conservation scenarios. We do not intend these results to provide a ranking of threats to guide conservation but rather showcase the complexity of setting global threat priorities and the importance of both accounting for the impact or frequency of threats.

Most importantly, our results show that to achieve global non-negative vertebrate population trends, we need to mitigate the effects of multiple threats (Figure 4 and S9). These findings offer evidence of ‘death by a thousand cuts’^36^, where multiple threats, over any one single feature are predictive of biodiversity declines. This is significant, as it implies that to reverse current vertebrate declines, conservation actions and management must target multiple threats simultaneously. For instance, according to our counterfactual tests, the only actions that could reverse global vertebrate declines would be mitigating habitat loss, climate change, and exploitation, or habitat loss, invasive species and exploitation at the same time (Figure S15). Managing these threats is challenging, given their direct links with the development of human societies^37–39^. Therefore, halting vertebrate declines requires ambitious conservation actions tackling multiple drivers of biodiversity loss^11,40^.

## Conclusions

There is much polarising debate as to whether wildlife populations are truly in decline^26,41^, with recent studies suggesting no net loss in vertebrate populations^17,42^, and that conservation efforts have been positive for biodiversity^25^. Here, using population-specific threat data and globally distributed time-series, we demonstrate that most vertebrate populations exposed to single and multiple threats are in decline across most systems and taxa. For instance, most vertebrate populations exposed to no threats show positive trends, while those exposed to at least one threat are either declining or show no net change (Figure 2; Table S1). These results highlight the importance of accounting for the impacts of multiple threats on populations to avoid underestimating the magnitude of biodiversity loss^19^.

Our analyses are based on the LPD, which is collected from peer-reviewed literature, grey literature, online databases, and data holders^26^, so these data come with inherent biases that need to be considered when interpreting global trends. Of the 25,054 population time-series in the LPD only 7,827 have information on threats, and only 3,129 were monitored for 10 years or more with at least 5 years of data. Our analyses mitigate these biases in our data by accounting for the temporal and spatial correlative effects, as well as disaggregating models by system and taxa. It is also important to note that our data does not include the strength and/or recurrency of the threats, which can also have a strong influence on populations trends^43^, and limits previous work that focus solely on qualitative changes in IUCN classifications (e.g. ^6^,^16^).

While we could not account for these different properties of threats, we quantify local population trends and how threats could explain the wide variety of trajectories observed in our study. For example, the differences between models fit to time series monitored for 10 and 20 years of data (Figure S10) may result from the predominantly seabird and fishery representation (Figure S11). These populations often display varying threat strengths through time due to fishing effort changes, which in turn influences time series dynamics^44,45^. Such non-linear dynamics can drive the overall linear estimate of threats to zero^46^. Future research targeting the impacts of the different strength and recurrence of threats on populations will be crucial to further understand the drivers of population declines^47,48^ and will help to develop more accurate biodiversity predictions^49^.

Overall, our results highlight that: (1) understanding the impacts of threats on the population dynamics of species provides key knowledge to fully characterise and predict the impacts of global change on biodiversity; and (2) mitigating multiple global threats should remain a conservation priority to halt biodiversity loss. Because of the limited resources allocated to conservation, rankings of the frequency of global threats are often used to prioritise conservation actions^1–3^. However, we show that the main threats impacting vertebrate populations are context-dependent^50,51^, and that accounting for both the impacts and frequency of threats is crucial to develop effective management actions. Particularly, multiple threats interacting at local scale can accelerate biodiversity loss and focusing efforts on mitigation of single threats might not be enough to halt the population declines.

## Materials and Methods

### Data

To measure the association between population trends and the different threats, both in isolation and with other stressors, we used one of the largest global population monitoring databases currently available, the Living Planet Database (LPD^26^). The LPD includes 25,054 population time-series of 4,392 species, each time-series has repeated monitoring surveys of the population abundance in a given area. These data are collated from a variety of sources, primarily from peer-reviewed literature, but also from grey literature, online databases, and data holders^26^. This information was digitised by the Living Planet Database team, following a standardised protocol (see details below).

Data sources containing population time-series for vertebrate species were collated by either scanning articles from journal issues within the conservation biology, wildlife management and ecology disciplines (this yielded most data sources); conducting keyword searches within academic and generic search engines; or contacting individual data holders to share data directly with the team (this yielded the least data sources). Data sources were selected if they met the criteria for inclusion in the LPD: single vertebrate species abundance estimates from a multi-annual survey, monitored using a consistent method and effort^52^ in the same location. Supporting texts in the publication, especially sections on limitations, were read to ensure that the abundance estimates were reliable and not an artifact, for example an increasing trend due to increased effort would not be included unless a correction factor was applied. Ancillary information on the survey location, ecology, and human activity (conservation interventions and threats) was collected from the data source alongside the abundance data. To ensure consistency, trained personnel recorded information from the original data sources using a set of guidelines. Once a person had been trained, all data entered are subject to an additional check by an experienced member of the LPD team to ensure quality control and promote consistency in decisions made.

Of the 25,054 population time-series making up the LPD (including confidential records), 7,827 population time-series contain information on whether the populations were exposed to threats. Based on information from the data source, for each publication it was first identified whether the population was threatened, not threatened, or whether its threat status was unknown. Threats were identified as direct or indirect human activities or processes that impacted the populations for at least 50% of the surveyed years, according to the original source of the time-series. Information on threat severity was not recorded as it was rarely available, and any mention of threats that was speculative in nature or only identified potential threats was not included. If the population was threatened, the number of threats the population was exposed to was recorded, from one to three. The information within the data sources was sometimes quantitative, e.g. stating number of individuals hunted annually, but most often it was reported in a qualitative way, e.g. describing a general pattern of hunting that impacts the populations. For this reason, and because the impact of the threat was rarely quantified in the data sources, broad categories describing the threat to the population were recorded: climate change, invasive species, habitat loss/degradation, exploitation, pollution, and diseases, following the Red List threat classification (Threats authority file Version 2.1; IUCN 2006a). As a guide, the subdivisions of the Red List threat classification were used to assign a threat to a broad category. We decided to use these broad categories to be able to compare the results of our study with those used in previous global biodiversity threat assessments^1,32^. Where there was more than one category option for a threat, the threat that better described the impact on the population was selected. For example, for amphibians we would list the threat of chytridiomycosis as a disease rather than as an invasive species (due to *Batrachochytrium dendrobatidis*). For species such as *Panthera tigris* which are threatened by a loss of prey base from hunting or competition with domestic species, we considered that habitat loss.

To ensure that our data could be associated with the potential impacts of threats on population trends, we only included time-series containing information on whether the population was exposed to threats at the time of the study (according to the original source). The number of threats populations were exposed to range from zero to three, as populations exposed to four or more threats were rarely found in the literature. To have sufficient information to appropriately capture the directional trends in abundance, we only included populations with at least ten years of monitoring data and a minimum of 5 data points^18,54^. We tested the implications of this choice of data quality by running additional analyses with ≥10 and ≥20 years of data. The influence of threats tended towards zero as time-series length increased (Figure S10), with the coverage of threats, systems and taxa strongly affected (Figure S11). This is likely due to threats and disturbances not acting linearly through time^55^, leading to a net zero linear trend^46^. The longest time-series were also heavily biased towards marine birds and fish, where ∼33% of time series are unthreatened in our dataset (Figure S11). We therefore present the results of the analyses run on data with ≥10 years and with a minimum of 5 data points to minimise system or taxa biases and include time-series with persistent threat effects.

Overall, our selection process resulted in 3,129 time-series of 1,281 species, including amphibians (83), birds (1195), fishes and elasmobranchs (947), mammals (698), and reptiles (206). These time-series covered all the continents (Figure 1), and freshwater (616), marine (1271), and terrestrial (1242) systems. The duration of data sets varied between 10 and 65 years, with a median duration of 25 years (Figure 1), covering a period between 1950 and 2019. Although the original dataset included data up to 2020, none of our selected time-series reached that year.

### Estimates of population trends

To estimate population trends, we fit a multilevel Bayesian linear model between the natural logarithm of abundance and year, and allowed year to interact with each threat/possible combination of threats. To identify the association between abundance and independent/interacting threats, we coded threats and threat combinations as binary factors. This resulted in a model matrix consisting of 36 columns representing the unique combinations of climate change, invasive species, habitat loss/degradation, exploitation, pollution, and disease. For example, if a population experienced exploitation and disease but not pollution, its model matrix would be: *exploitation = 1, disease = 1, pollution = 0, exploitation.disease = 1, exploitation.pollution = 0, disease.pollution = 0, exploitation.disease.pollution = 0*. We consequently can decompose the additive effects of exploitation and disease (*exploitation = 1, disease = 1*) from their interaction (*exploitation.disease = 1*). Defining threats in this way is necessitated by the qualitative threat categorisation described above and allows counterfactual analyses.

To account for temporal non-independence, we modelled the population level time-series with a discrete autoregressive-1 (ar1) temporal process, which assumes neighbouring abundance observations within a time-series will be more similar. Further, to capture spatial and phylogenetic correlative non-independence, we fit covariance structures between species identities and sites^27^. Phylogeny was modelled via uncorrelated slopes between species and populations, while shared correlations between sites were fit via a shared slope covariance structure. This site covariance matrix was derived using the Haversine (spherical) distance between each site. To improve run time in the largest datasets we rounded site coordinates to the nearest integer i.e. a latitude of 10.65 was set to 11. We normalised this spatial matrix between 0 and 1, with values close to 1 indicating neighbouring sites, whilst values approaching 0 indicate distant sites.

Year and abundance were centred at zero by subtracting the mean time series year/abundance from each time series’ year/abundance. This parameterization effectively fixes intercepts at 0 for each slope and accounts for random intercept variation without increasing the number of estimated parameters. Consequently, no random intercepts were necessary.

The final model structure was as follows:

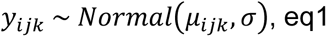

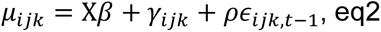

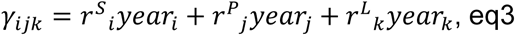

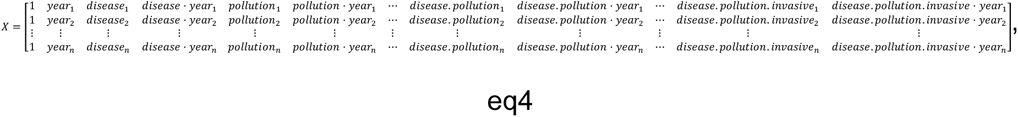

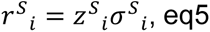

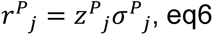

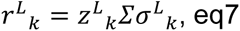

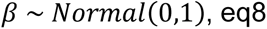

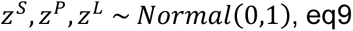

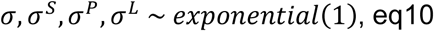

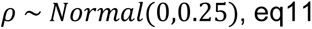

Specifically, we assumed abundance (*y*) is a gaussian normal distributed variable with mean *μ* and variance *σ,* consisting of *n* observations. *X* represents the fixed effects model matrix containing year and threat variables, with *β* the associated fixed effect coefficients. *γ* is therefore the random effect coefficients which are controlled by the independent random slopes for species (*r^S^*, index *i*) and population (*r^P^*, index *j*), and the correlated site (*r^L^*, index *k*) random slopes. Due to challenges fitting the random hyperparameters, the model was reparametrized in to a “non-centred” form to prevent correlations between hyperparameters^56^. Consequently, rather than modelling slopes varying around the overall year coefficient, we introduce a standard normal parameter for each random effect (*z^S^, z^P^, z^L^*) which is then multiplied by each random effect’s respective sigma hyperprior (*σ^S^*, *σ^P^*, *σ^L^*). *r^L^* (the correlated site slopes) is also dependent upon the additional variance-covariance *Σ* which specifies that covariance is present in neighbouring sites. *Σ* is populated by the Haversine distances between each site. Finally, non-centred parameterisation allows temporal non-independence to be modelled by introducing the correlation (*ρ*) between neighbouring residuals (*ε*) in the linear predictor rather than the observational error term (*σ*) as is commonly parameterised.

Further, we specify that all time-series, and in turn species, share the same autocorrelation parameter (*ρ*). This model structure was then fit across subsets of the LPI dataset to disentangle differential responses of taxa and ecosystems to threats. First, a global model was fit across all appropriate time series to give insight into general trends and relationships between abundance and threats. Second, system level models were fit upon freshwater, marine, and terrestrial time series alone. Finally, taxa specific models were fit upon time series classified as amphibians, birds, fishes, mammals, or reptiles. The category “fishes” incorporated the taxonomic groups Holocephali, Elasmobranchii, Myxini, Cephalaspidomorphi, Actinopterygii, and Sarcopterygii. It was necessary to fit models by system and/or taxa due to the large number of parameters estimated in each model. For example, the inclusion of a taxa variable with five levels (amphibian, bird, fish, mammal, and reptile) would increase the number of fixed effect coefficients in the global model from 74 (the 36 threat combinations, year, the interaction between year and each threat combination, and global intercept) to 370. Model structure was identical across models with the spatial variance-covariance matrix *Σ* recalculated for each data subset.

All models were fit using the brms package v2.22.0^57^ in R v4.4.0^58^. Models were run for 5000 iterations over 4 chains, with a warmup of 2500 iterations. This resulted in a total of 20000 draws per model. All distributions reported in this study were constructed from 1000 random draws of the posterior distribution.

### Multilevel Bayesian models diagnostics

To check the validity of our multilevel Bayesian models we ran a set of diagnostics. We inspected model convergence by visually examining trace plots and using Rhat values (the ratio of the effective sample size to the overall number of iterations, with values close to one indicating convergence).

Furthermore, we evaluated the model fit by exploring the distribution of the residuals, their variance, their autocorrelation, and the posterior predictive checks (Figures S12-S15). When the model fits the data, we would expect the residuals to follow a Gaussian distribution and to show constant variance, as showed in our models (Figures S12-S135). In addition, the posterior predictive checks compare the distribution of the data with the predictions from the model, so if the model is well fitted, the predictions should overlap the data, as showed in our models (Figure S14). Finally, our models displayed no evidence of autocorrelation in their residuals (Figure S15)

### Influence of threats on population trends

To quantify the trends of populations exposed to single and multiple threats on population trends, we examined coefficients and extracted predictions made by the fitted multilevel Bayesian models. As each threat factor was coded as binary, the year coefficient represents trends in the absence of threats, and each threat/threat interaction coefficient is therefore the moderation of this no-threat trend.

Additionally, to estimate the ultimate association of threats with population trends, we estimated the conditional effect for each combination of threats for each year contained within the LPD. This generates a predictive posterior distribution of year-on-year log abundances which we then converted to trend by taking the mean derivative (a.k.a. the slope) using the equation 12 on a draw-by-draw basis:

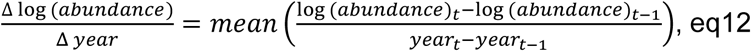

Henceforth, the term “trend” refers to the derivative of the predicted slope rather than the raw coefficients.

The inclusion of both random and fixed effects in our analysis has the potential to confound predicted trends. We therefore validated our predictions by comparing the variability of trends predicted using random effects alone versus those made with the inclusion of threat fixed effects. To achieve this, we generated null model predictions by estimating the conditional trends (i.e. random effects set to 0) for populations where all threats were “removed” and set to 0. The null trend posterior was then compared to two separate scenarios estimating the contribution of random vs fixed effects to the observed trends. Random effect predictions were made upon the null dataset but included all random effect identities (species, population, site etc). We then averaged these predictions across all time series draws to estimate a marginal trend posterior. Conversely, fixed effect contributions were estimated via conditional trends with all threats “present” and set to 1. Each of these predictions were then differenced from the null model posterior to create a distribution of trend “contributions”. If the trend difference is zero, then that effect is not contributing to the direction of trend. We anticipate that random effects will not influence the direction of trends but increase the uncertainty of estimates^27^.

### Interactive vs additive effects of threats

We also classified the form of interactive threat effects in a separate analysis using the above-mentioned multilevel Bayesian model. By including or excluding threat data, we estimated conditional trends given independent and/or interacting threats and used the difference in trends to classify interaction type. In our model, the additive contribution of threats to trend can be predicted by including solely the independent threat columns in the threat model matrix. For example, if we were interested in the additive effect of exploitation and disease, we would code the model matrix as: *exploitation = 1, disease = 1, pollution = 0, exploitation.disease = 0, exploitation.pollution = 0, disease.pollution = 0, exploitation.disease.pollution = 0*. Exploitation and disease are therefore the only contributing threats to the conditional trend predictions, with their interactions not considered. Consequently, we can alter the model matrix to include exploitation and disease’s interaction by “turning on” the interactive column: *exploitation = 1, disease = 1, pollution = 0, exploitation.disease = 1, exploitation.pollution = 0, disease.pollution = 0, exploitation.disease.pollution = 0*. We therefore have two posteriors representing the additive and interactive contributions of threats to population trend.

There is still an ongoing debate on the best approaches to classify interactive effects of multiple threats^23,59,60^. In this study we considered that the null model was the simple additive effect (sum of the individual effects), because it was the most conservative approach based on the scale of our study^60^. We included organisms from different taxonomic groups and ecosystems, and given we lacked information about the mechanistic effects of each threat, we decided to maintain a simple additive null effect^60^. Consequently, we differenced the interacting threat posterior from the additive null and used the direction of difference to classify threat contributions as additive, antagonistic or synergistic (Figure S1). If the 80% credible interval for the differenced posterior transgressed zero, then the threat combination was classified as additive. We assume that threats contribute negatively to trends, and so, if the difference in trend is negative, we classified that threat combination as synergistic – the interaction of threats generate a trend posterior more negative than the additive threat posterior. Conversely, if the difference in trend is positive, we classified that threat combination as antagonistic as the interacting threat posterior is more positive than the additive equivalent. See Figure S1 for a visualisation of this process.

It is worth mentioning that our approach to quantify the interactive effects assumes that the population trends measured are predicted by the threats. Nevertheless, in this study we did not perform any causal analysis, which could establish causal links between the threats and the observed population trends. It is possible that other confounding factors may have played a role in shaping these population trends, but it was out of the scope of this manuscript to establish these causal links. As a result, we advise caution when interpreting the results derived from our analyses of interactive effects, given that we assume causal links that were not formally tested.

### Counterfactual scenarios

To explore what would happen to vertebrate population predictions if threat contributions were removed, we developed six counterfactual scenarios. Each counterfactual scenario represents the predicted population trends where the potential effect of the different single threats (exploitation, climate change, habitat loss, disease, invasive species and pollution) was removed from the populations exposed to any number of threats. To achieve this, we took the following steps:

1. We first dropped all populations in our dataset that were unthreatened.
2. We then modified the model matrix for each threat in turn, by setting all columns where that threat is present to 0. For example, a counterfactual removing exploitation from a population experiencing exploitation and disease would be coded as: *exploitation = 0, disease = 1, pollution = 0, exploitation.disease = 0, exploitation.pollution = 0, disease.pollution = 0, exploitation.disease.pollution = 0* (exploitation is removed, disease remains, but the interaction is also removed).
3. We then estimated the trend for each population given the counterfactual model matrix to create a distribution of population trends given the individual removal of each threat.
4. We finally compared the counterfactual distribution to the mean trend of the unmodified model matrix to identify whether removal of the threat resulted in more positive predicted trends. The difference is reported as the percentage increase/decrease in trend using the equation 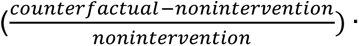 100 where *counterfactual* is the estimated trend for hypothetical populations where a threat is removed and *non-intervention* represents the estimated trend in the absence of any intervention. We repeated this procedure for each single threat.

## Acknowledgements.

PC is supported by Leverhulme grant RPG-2019-368. L.M. is funded by WWF UK and WWF Netherlands.

## Author contributions

P.C., L.M. and C.C. conceived the idea with inputs from all the authors. P.C., D.O, T.J., and C.C. performed the analyses. L.M., R.F., and V.M. collected the data. P.C., D.O, L.M., and C.C. wrote the manuscript with contributions from all authors.

## Competing interests

The authors declare no competing interests.

Correspondence and requests for materials should be addressed to.

**Supplementary information** is available for this paper.

## Statement of data availability

The data was obtained from the https://livingplanetindex.org/data_portal. The population trends data will be deposited in Zotero upon manuscript acceptance.

## Statement of code availability

R code will be deposited in Zotero upon manuscript acceptance. During the reviewing process the code will be available at https://github.com/duncanobrien/multiple-threats.

