## Supplementary material for "Halting predicted vertebrate declines requires tackling multiple drivers of biodiversity loss": Figure S

1. School of Biological Sciences, University of Bristol, Bristol, UK
2. Departament de Biologia Evolutiva, Ecologia i Ciències Ambientals, Universitat de Barcelona, Barcelona, Spain
3. Institut de Recerca de la Biodiversitat, Universitat de Barcelona (UB), Barcelona, Spain
4. Institute of Zoology, Zoological Society of London, London, UK
5. School of Biosciences, University of Sheffield, Sheffield, UK
* These authors contributed equally to this work
† These authors contributed equally to this work

#### Contents

**1 Appendix S1: Extended methods** **2**

**2 Appendix S2: Extended results** **3**

**3 Appendix S3: Model checks** **28**

### 1 Appendix S1: Extended methods

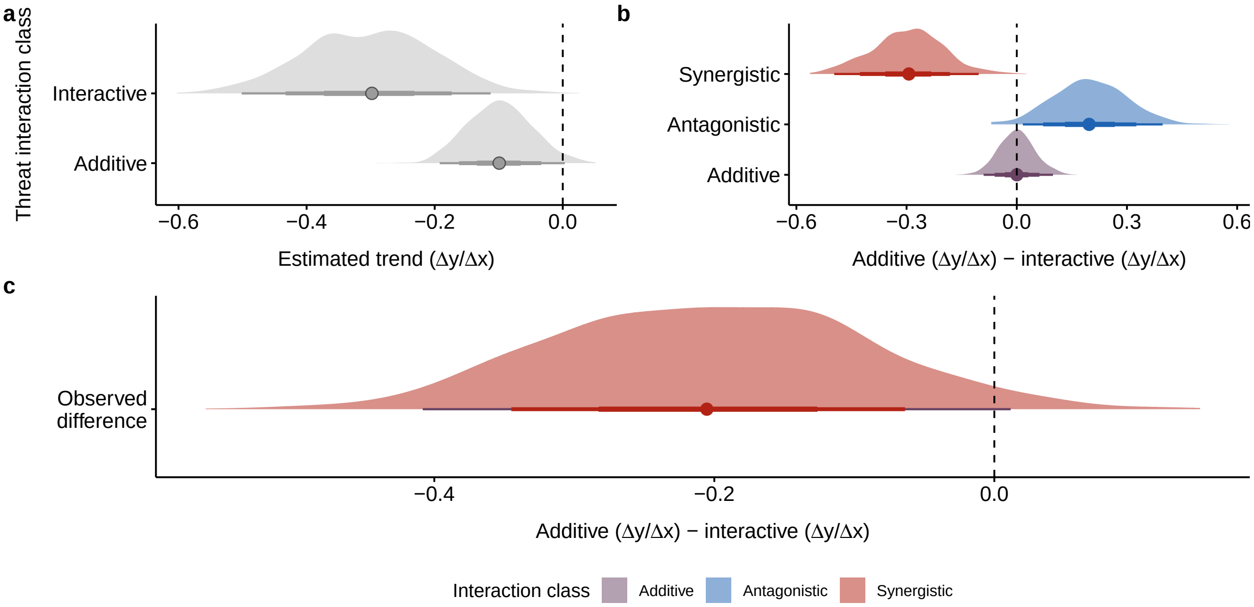


Figure S1: Demonstration of the threat classification protocol. (a) Trends are first predicted for each threat in additive or interactive combinations. (b) If the difference between predicted interactive trends and additive trends is calculated, there are three possible classes: additive (no difference predictions made by additive and interacting threats), antagonistic (interacting threat predictions are less negative than additive predictions), or synergistic (interacting threat predictions are more negative than additive predictions). Classes are defined using the 80% credible interval. We consider that a synergy 'worsens' trends - i.e. becomes more negative. (c) The actual difference between predictions made in panel a.

### 2 Appendix S2: Extended results

Table S1: Model coeffcients for global population trends. Median represents the median of the posterior distribution. CI low and high are the lower and higher values of the 95% credible interval. Rhat is the ratio of the effective sample size to the overall number of iterations, with values close to one indicating convergence values

| Parameter | Median | CI_low | CI_high | Rhat | ESS |
|---|---|---|---|---|---|
| Intercept | -0.002 | -0.011 | 0.008 | 1 | 16766.77 |
| scaled_year | 0.015 | -0.154 | 0.180 | 1 | 5239.48 |
| pollution1 | 0.002 | -0.076 | 0.078 | 1 | 6679.82 |
| habitatl1 | -0.002 | -0.025 | 0.021 | 1 | 11880.81 |
| climatechange1 | -0.006 | -0.053 | 0.043 | 1 | 10845.48 |
| invasive1 | 0.001 | -0.052 | 0.054 | 1 | 9582.01 |
| exploitation1 | -0.002 | -0.018 | 0.015 | 1 | 15430.68 |
| disease1 | 0.003 | -0.078 | 0.081 | 1 | 8577.92 |
| pollution.habitatl1 | 0.002 | -0.089 | 0.093 | 1 | 6692.33 |
| pollution.climatechange1 | 0.011 | -1.012 | 1.030 | 1 | 5600.75 |
| pollution.invasive1 | 0.023 | -1.019 | 1.055 | 1 | 5762.86 |
| pollution.exploitation1 | -0.018 | -0.112 | 0.078 | 1 | 6942.95 |
| pollution.disease1 | -0.014 | -0.263 | 0.239 | 1 | 8122.48 |
| habitatl.climatechange1 | 0.007 | -0.067 | 0.076 | 1 | 9522.42 |
| habitatl.invasive1 | 0.002 | -0.076 | 0.078 | 1 | 8657.07 |
| habitatl.exploitation1 | 0.001 | -0.039 | 0.042 | 1 | 11701.48 |
| habitatl.disease1 | 0.008 | -0.112 | 0.129 | 1 | 8171.24 |
| climatechange.invasive1 | -0.002 | -0.295 | 0.299 | 1 | 8996.20 |
| climatechange.exploitation1 | 0.010 | -0.109 | 0.130 | 1 | 10428.10 |
| climatechange.disease1 | -0.009 | -0.295 | 0.275 | 1 | 11005.17 |
| invasive.exploitation1 | -0.005 | -0.101 | 0.096 | 1 | 9620.00 |
| invasive.disease1 | -0.037 | -0.226 | 0.153 | 1 | 8536.11 |
| exploitation.disease1 | 0.001 | -0.192 | 0.196 | 1 | 8136.06 |
| pollution.habitatl.climatechange1 | -0.010 | -1.043 | 1.019 | 1 | 5592.53 |
| pollution.habitatl.invasive1 | -0.023 | -1.062 | 1.030 | 1 | 5684.72 |
| pollution.habitatl.exploitation1 | 0.014 | -0.110 | 0.141 | 1 | 6989.64 |
| pollution.habitatl.disease1 | -0.005 | -0.288 | 0.291 | 1 | 7668.28 |
| pollution.climatechange.invasive1 | -0.025 | -1.299 | 1.246 | 1 | 6833.08 |
| pollution.climatechange.disease1 | 0.039 | -1.000 | 1.112 | 1 | 5780.83 |
| pollution.invasive.exploitation1 | 0.027 | -1.003 | 1.082 | 1 | 5794.27 |
| pollution.exploitation.disease1 | 0.035 | -0.313 | 0.382 | 1 | 7420.34 |
| habitatl.climatechange.invasive1 | 0.015 | -0.328 | 0.356 | 1 | 8993.30 |
| habitatl.climatechange.exploitation1 | -0.019 | -0.169 | 0.133 | 1 | 8989.49 |
| habitatl.climatechange.disease1 | 0.002 | -0.338 | 0.339 | 1 | 10009.82 |
| habitatl.invasive.exploitation1 | 0.002 | -0.161 | 0.162 | 1 | 10466.64 |
| habitatl.invasive.disease1 | 0.030 | -0.266 | 0.326 | 1 | 9265.35 |
| habitatl.exploitation.disease1 | -0.013 | -0.273 | 0.250 | 1 | 8028.88 |
| climatechange.invasive.exploitation1 | 0.006 | -0.435 | 0.446 | 1 | 9521.38 |
| invasive.exploitation.disease1 | 0.013 | -0.359 | 0.388 | 1 | 9400.91 |
| scaled_year:pollution1 | -0.034 | -0.068 | 0.001 | 1 | 3295.92 |
| scaled_year:habitatl1 | -0.021 | -0.032 | -0.011 | 1 | 5386.22 |
| scaled_year:climatechange1 | -0.047 | -0.068 | -0.025 | 1 | 4718.61 |
| scaled_year:invasive1 | -0.075 | -0.101 | -0.049 | 1 | 5220.89 |
| scaled_year:exploitation1 | -0.026 | -0.035 | -0.017 | 1 | 5244.63 |
| scaled_year:disease1 | -0.062 | -0.097 | -0.028 | 1 | 4449.70 |
| scaled_year:pollution.habitatl1 | 0.044 | 0.004 | 0.085 | 1 | 3453.45 |
| scaled_year:pollution.climatechange1 | -0.016 | -1.039 | 1.002 | 1 | 5746.41 |
| scaled_year:pollution.invasive1 | 0.013 | -0.967 | 1.033 | 1 | 5182.89 |
| scaled_year:pollution.exploitation1 | 0.040 | -0.003 | 0.082 | 1 | 3344.74 |
| scaled_year:pollution.disease1 | 0.011 | -0.104 | 0.128 | 1 | 4235.49 |
| scaled_year:habitatl.climatechange1 | 0.040 | 0.009 | 0.071 | 1 | 4264.91 |
| scaled_year:habitatl.invasive1 | 0.074 | 0.034 | 0.114 | 1 | 4961.77 |
| scaled_year:habitatl.exploitation1 | 0.009 | -0.010 | 0.027 | 1 | 4138.91 |
| scaled_year:habitatl.disease1 | 0.084 | 0.024 | 0.142 | 1 | 4779.72 |

Table S1: Model coefficents for global population trends. Median represents the median of the posterior distribution. CI low and high are the lower and higher values of the 95% credible interval. Rhat is the ratio of the effective sample size to the overall number of iterations, with values close to one indicating convergence values *(continued)*

| Parameter | Median | CI_low | CI_high | Rhat | ESS |
|---|---|---|---|---|---|
| scaled_year:climatechange.invasive1 | 0.090 | -0.058 | 0.235 | 1 | 6518.40 |
| scaled_year:climatechange.exploitation1 | 0.055 | 0.006 | 0.104 | 1 | 6193.38 |
| scaled_year:climatechange.disease1 | 0.112 | -0.014 | 0.235 | 1 | 7038.99 |
| scaled_year:invasive.exploitation1 | 0.121 | 0.075 | 0.167 | 1 | 5631.31 |
| scaled_year:invasive.disease1 | 0.115 | 0.035 | 0.196 | 1 | 5166.76 |
| scaled_year:exploitation.disease1 | 0.012 | -0.071 | 0.096 | 1 | 4328.87 |
| scaled_year:pollution.habitatl.climatechange1 | -0.017 | -1.038 | 1.008 | 1 | 5769.50 |
| scaled_year:pollution.habitatl.invasive1 | -0.041 | -1.065 | 0.952 | 1 | 5204.00 |
| scaled_year:pollution.habitatl.exploitation1 | -0.070 | -0.126 | -0.015 | 1 | 3742.15 |
| scaled_year:pollution.habitatl.disease1 | -0.041 | -0.173 | 0.092 | 1 | 4344.52 |
| scaled_year:pollution.climatechange.invasive1 | 0.154 | -1.097 | 1.393 | 1 | 6081.11 |
| scaled_year:pollution.climatechange.disease1 | -0.159 | -1.167 | 0.869 | 1 | 5775.97 |
| scaled_year:pollution.invasive.exploitation1 | -0.130 | -1.155 | 0.860 | 1 | 5181.62 |
| scaled_year:pollution.exploitation.disease1 | 0.083 | -0.077 | 0.235 | 1 | 4300.89 |
| scaled_year:habitatl.climatechange.invasive1 | -0.070 | -0.229 | 0.099 | 1 | 6275.61 |
| scaled_year:habitatl.climatechange.exploitation1 | -0.031 | -0.097 | 0.037 | 1 | 5321.56 |
| scaled_year:habitatl.climatechange.disease1 | -0.166 | -0.317 | -0.018 | 1 | 6375.00 |
| scaled_year:habitatl.invasive.exploitation1 | -0.133 | -0.207 | -0.059 | 1 | 5567.15 |
| scaled_year:habitatl.invasive.disease1 | -0.152 | -0.278 | -0.027 | 1 | 5514.69 |
| scaled_year:habitatl.exploitation.disease1 | 0.005 | -0.106 | 0.116 | 1 | 4259.27 |
| scaled_year:climatechange.invasive.exploitation1 | -0.109 | -0.322 | 0.103 | 1 | 7490.15 |
| scaled_year:invasive.exploitation.disease1 | -0.210 | -0.365 | -0.057 | 1 | 5027.81 |

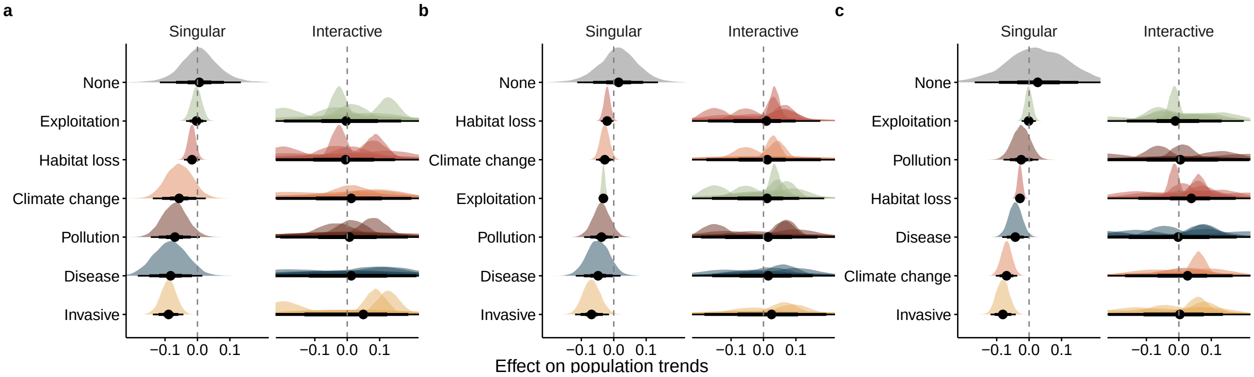


Figure S2: Effects of threats on vertebrate population trends across realms. 95% credible intervals of the effects of single and interacting threats upon (a) freshwater, (b) marine, and (c) terrestrial time series trends. The dashed vertical line shows zero influence – i.e., no effect of the factors – while the None parameter is the trend in the absence of threats. The remaining parameters represent modifications of this None trend. The credible intervals are based on 1,000 samples from the posterior distribution of the model coefficients.

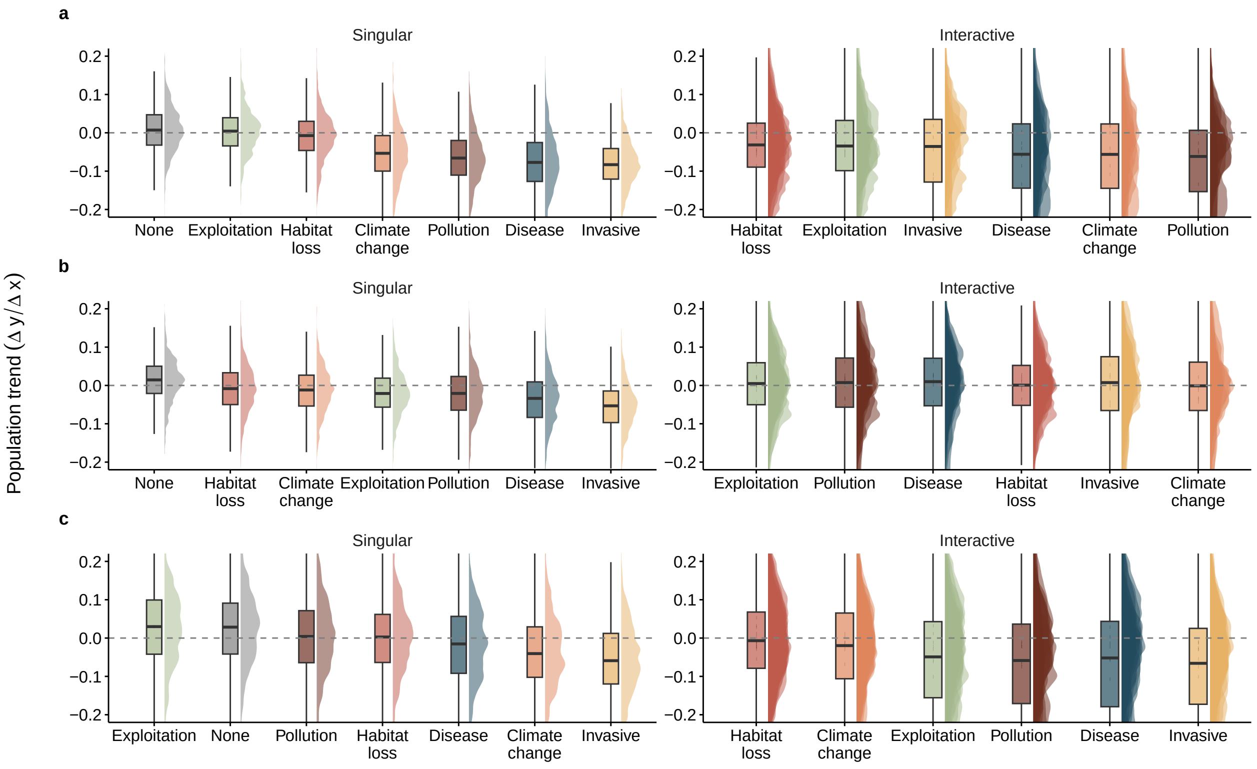


Figure S3: Ultimate effect of threats on vertebrate population trends across realms. 95% credible intervals of the ultimate estimated trend in the presence of single and interacting threats for (a) freshwater, (b) marine, and (c) terrestrial time series. The dashed horizontal line shows the zero-slope – i.e., no effect of the factors. The credible intervals are based on 1,000 samples from the posterior distribution of the model predictions where threats are included/excluded from the model matrix.

Table S2: Model coefficents for global population trends across systems. Median represents the median of the posterior distribution. CI low and high are the lower and higher values of the 95% credible interval. Rhat is the ratio of the effective sample size to the overall number of iterations, with values close to one indicating convergence values.

| System | Parameter | Median | CI_low | CI_high | Rhat | ESS |
|---|---|---|---|---|---|---|
| Freshwater | Intercept | -0.004 | -0.030 | 0.022 | 1 | 12754.95 |
| Freshwater | scaled_year | 0.006 | -0.116 | 0.134 | 1 | 5521.06 |
| Freshwater | pollution1 | -0.010 | -0.143 | 0.124 | 1 | 7146.66 |
| Freshwater | habitatl1 | 0.001 | -0.045 | 0.048 | 1 | 12127.95 |
| Freshwater | climatechange1 | -0.010 | -0.174 | 0.153 | 1 | 10972.94 |
| Freshwater | invasive1 | 0.009 | -0.091 | 0.103 | 1 | 8280.23 |
| Freshwater | exploitation1 | -0.001 | -0.060 | 0.060 | 1 | 11073.57 |
| Freshwater | disease1 | -0.012 | -0.209 | 0.180 | 1 | 12865.39 |
| Freshwater | pollution.habitatl1 | 0.013 | -0.136 | 0.163 | 1 | 7589.37 |
| Freshwater | pollution.climatechange1 | 0.014 | -0.391 | 0.426 | 1 | 13203.35 |
| Freshwater | pollution.invasive1 | -0.002 | -0.373 | 0.358 | 1 | 10973.76 |
| Freshwater | pollution.exploitation1 | -0.005 | -0.182 | 0.167 | 1 | 7431.10 |
| Freshwater | pollution.disease1 | 0.023 | -0.297 | 0.342 | 1 | 9581.91 |
| Freshwater | habitatl.climatechange1 | -0.001 | -0.232 | 0.228 | 1 | 10561.24 |
| Freshwater | habitatl.invasive1 | 0.001 | -0.119 | 0.123 | 1 | 7862.94 |
| Freshwater | habitatl.exploitation1 | 0.003 | -0.090 | 0.099 | 1 | 10051.42 |
| Freshwater | habitatl.disease1 | 0.008 | -0.267 | 0.284 | 1 | 10721.85 |
| Freshwater | climatechange.invasive1 | -0.003 | -0.393 | 0.380 | 1 | 11628.41 |
| Freshwater | climatechange.exploitation1 | 0.018 | -0.302 | 0.342 | 1 | 12398.20 |
| Freshwater | climatechange.disease1 | 0.013 | -0.267 | 0.297 | 1 | 11557.64 |
| Freshwater | invasive.exploitation1 | -0.009 | -0.145 | 0.127 | 1 | 8448.46 |
| Freshwater | exploitation.disease1 | 0.016 | -0.327 | 0.365 | 1 | 10966.37 |
| Freshwater | pollution.habitatl.invasive1 | -0.004 | -0.367 | 0.373 | 1 | 11072.99 |
| Freshwater | pollution.habitatl.exploitation1 | 0.003 | -0.202 | 0.215 | 1 | 7960.96 |
| Freshwater | pollution.habitatl.disease1 | -0.016 | -0.377 | 0.342 | 1 | 10667.78 |
| Freshwater | pollution.climatechange.disease1 | 0.011 | -0.407 | 0.419 | 1 | 11130.40 |
| Freshwater | pollution.exploitation.disease1 | 0.009 | -0.390 | 0.404 | 1 | 9905.53 |
| Freshwater | habitatl.climatechange.invasive1 | -0.010 | -0.396 | 0.366 | 1 | 11761.48 |
| Freshwater | habitatl.climatechange.exploitation1 | 0.003 | -0.349 | 0.345 | 1 | 11612.46 |
| Freshwater | habitatl.climatechange.disease1 | -0.002 | -0.336 | 0.344 | 1 | 10364.05 |
| Freshwater | habitatl.invasive.exploitation1 | 0.000 | -0.239 | 0.239 | 1 | 10506.96 |
| Freshwater | habitatl.exploitation.disease1 | -0.007 | -0.399 | 0.377 | 1 | 12460.22 |
| Freshwater | scaled_year:pollution1 | -0.069 | -0.144 | 0.007 | 1 | 3654.67 |
| Freshwater | scaled_year:habitatl1 | -0.017 | -0.042 | 0.007 | 1 | 2631.19 |
| Freshwater | scaled_year:climatechange1 | -0.056 | -0.138 | 0.026 | 1 | 5280.14 |
| Freshwater | scaled_year:invasive1 | -0.089 | -0.137 | -0.043 | 1 | 3431.43 |
| Freshwater | scaled_year:exploitation1 | -0.003 | -0.035 | 0.028 | 1 | 3903.79 |
| Freshwater | scaled_year:disease1 | -0.083 | -0.184 | 0.015 | 1 | 5968.31 |
| Freshwater | scaled_year:pollution.habitatl1 | 0.078 | -0.009 | 0.161 | 1 | 3466.55 |
| Freshwater | scaled_year:pollution.climatechange1 | -0.044 | -0.426 | 0.353 | 1 | 9896.34 |
| Freshwater | scaled_year:pollution.invasive1 | -0.025 | -0.373 | 0.326 | 1 | 10254.28 |
| Freshwater | scaled_year:pollution.exploitation1 | 0.015 | -0.080 | 0.112 | 1 | 3966.57 |
| Freshwater | scaled_year:pollution.disease1 | 0.013 | -0.272 | 0.301 | 1 | 7130.72 |
| Freshwater | scaled_year:habitatl.climatechange1 | 0.014 | -0.115 | 0.143 | 1 | 5577.90 |
| Freshwater | scaled_year:habitatl.invasive1 | 0.087 | 0.023 | 0.153 | 1 | 3644.98 |
| Freshwater | scaled_year:habitatl.exploitation1 | -0.026 | -0.073 | 0.022 | 1 | 3440.82 |
| Freshwater | scaled_year:habitatl.disease1 | 0.121 | -0.056 | 0.301 | 1 | 6356.47 |
| Freshwater | scaled_year:climatechange.invasive1 | 0.034 | -0.316 | 0.390 | 1 | 9025.21 |
| Freshwater | scaled_year:climatechange.exploitation1 | 0.071 | -0.126 | 0.269 | 1 | 7651.56 |
| Freshwater | scaled_year:climatechange.disease1 | 0.090 | -0.087 | 0.268 | 1 | 5993.53 |
| Freshwater | scaled_year:invasive.exploitation1 | 0.127 | 0.057 | 0.198 | 1 | 3487.54 |
| Freshwater | scaled_year:exploitation.disease1 | 0.014 | -0.313 | 0.337 | 1 | 7553.37 |
| Freshwater | scaled_year:pollution.habitatl.invasive1 | -0.017 | -0.371 | 0.338 | 1 | 10481.86 |
| Freshwater | scaled_year:pollution.habitatl.exploitation1 | -0.034 | -0.151 | 0.086 | 1 | 3636.54 |
| Freshwater | scaled_year:pollution.habitatl.disease1 | -0.041 | -0.361 | 0.267 | 1 | 6823.34 |
| Freshwater | scaled_year:pollution.climatechange.disease1 | -0.040 | -0.429 | 0.345 | 1 | 10929.37 |
| Freshwater | scaled_year:pollution.exploitation.disease1 | 0.086 | -0.281 | 0.451 | 1 | 8700.69 |

Table S2: Model coefficents for global population trends across systems. Median represents the median of the posterior distribution. CI low and high are the lower and higher values of the 95% credible interval. Rhat is the ratio of the effective sample size to the overall number of iterations, with values close to one indicating convergence values. *(continued)*

| System | Parameter | Median | CI_low | CI_high | Rhat | ESS |
|---|---|---|---|---|---|---|
| Freshwater | scaled__year:habitatl.climatechange.invasive1 | 0.040 | -0.317 | 0.392 | 1 | 9106.81 |
| Freshwater | scaled__year:habitatl.climatechange.exploitation1 | 0.008 | -0.228 | 0.241 | 1 | 6843.99 |
| Freshwater | scaled__year:habitatl.climatechange.disease1 | -0.164 | -0.408 | 0.080 | 1 | 6026.35 |
| Freshwater | scaled__year:habitatl.invasive.exploitation1 | -0.202 | -0.326 | -0.078 | 1 | 4507.62 |
| Freshwater | scaled__year:habitatl.exploitation.disease1 | -0.075 | -0.416 | 0.267 | 1 | 9330.24 |
| Marine | Intercept | 0.001 | -0.015 | 0.018 | 1 | 18797.44 |
| Marine | scaled__year | 0.015 | -0.113 | 0.136 | 1 | 6179.38 |
| Marine | pollution1 | 0.009 | -0.104 | 0.126 | 1 | 7315.41 |
| Marine | habitatl1 | -0.020 | -0.065 | 0.026 | 1 | 11427.51 |
| Marine | climatechange1 | -0.016 | -0.081 | 0.049 | 1 | 12419.27 |
| Marine | invasive1 | -0.041 | -0.168 | 0.086 | 1 | 15226.51 |
| Marine | exploitation1 | -0.007 | -0.030 | 0.016 | 1 | 17323.53 |
| Marine | disease1 | -0.002 | -0.171 | 0.170 | 1 | 10525.81 |
| Marine | pollution.habitatl1 | 0.005 | -0.128 | 0.140 | 1 | 6915.88 |
| Marine | pollution.climatechange1 | 0.004 | -0.342 | 0.335 | 1 | 10109.16 |
| Marine | pollution.invasive1 | 0.004 | -0.338 | 0.341 | 1 | 12127.98 |
| Marine | pollution.exploitation1 | -0.014 | -0.149 | 0.116 | 1 | 7317.14 |
| Marine | pollution.disease1 | -0.014 | -0.258 | 0.223 | 1 | 9785.11 |
| Marine | habitatl.climatechange1 | 0.031 | -0.072 | 0.136 | 1 | 10426.30 |
| Marine | habitatl.invasive1 | 0.027 | -0.165 | 0.219 | 1 | 14157.95 |
| Marine | habitatl.exploitation1 | 0.014 | -0.058 | 0.086 | 1 | 10208.42 |
| Marine | habitatl.disease1 | 0.026 | -0.166 | 0.214 | 1 | 9775.90 |
| Marine | climatechange.invasive1 | 0.017 | -0.288 | 0.328 | 1 | 12221.42 |
| Marine | climatechange.exploitation1 | 0.019 | -0.110 | 0.145 | 1 | 11887.43 |
| Marine | invasive.exploitation1 | 0.042 | -0.169 | 0.249 | 1 | 10915.28 |
| Marine | invasive.disease1 | 0.008 | -0.336 | 0.352 | 1 | 12886.71 |
| Marine | exploitation.disease1 | 0.007 | -0.250 | 0.269 | 1 | 11304.97 |
| Marine | pollution.habitatl.climatechange1 | -0.014 | -0.364 | 0.338 | 1 | 10969.60 |
| Marine | pollution.habitatl.exploitation1 | -0.001 | -0.178 | 0.173 | 1 | 7490.56 |
| Marine | pollution.habitatl.disease1 | -0.016 | -0.308 | 0.272 | 1 | 10275.71 |
| Marine | pollution.climatechange.invasive1 | 0.008 | -0.402 | 0.415 | 1 | 13669.66 |
| Marine | pollution.invasive.exploitation1 | -0.013 | -0.387 | 0.354 | 1 | 12165.17 |
| Marine | pollution.exploitation.disease1 | 0.006 | -0.314 | 0.330 | 1 | 11351.94 |
| Marine | habitatl.climatechange.invasive1 | 0.007 | -0.375 | 0.398 | 1 | 16826.52 |
| Marine | habitatl.climatechange.exploitation1 | -0.048 | -0.226 | 0.132 | 1 | 10347.15 |
| Marine | habitatl.invasive.exploitation1 | -0.003 | -0.277 | 0.271 | 1 | 10802.03 |
| Marine | habitatl.invasive.disease1 | 0.008 | -0.373 | 0.389 | 1 | 14973.33 |
| Marine | habitatl.exploitation.disease1 | -0.012 | -0.384 | 0.360 | 1 | 16424.58 |
| Marine | climatechange.invasive.exploitation1 | -0.009 | -0.375 | 0.352 | 1 | 13443.44 |
| Marine | invasive.exploitation.disease1 | 0.001 | -0.386 | 0.380 | 1 | 13258.15 |
| Marine | scaled__year:pollution1 | -0.038 | -0.091 | 0.014 | 1 | 3927.99 |
| Marine | scaled__year:habitatl1 | -0.020 | -0.040 | -0.001 | 1 | 4770.87 |
| Marine | scaled__year:climatechange1 | -0.028 | -0.055 | 0.000 | 1 | 4786.87 |
| Marine | scaled__year:invasive1 | -0.068 | -0.119 | -0.015 | 1 | 6076.94 |
| Marine | scaled__year:exploitation1 | -0.032 | -0.043 | -0.021 | 1 | 4280.51 |
| Marine | scaled__year:disease1 | -0.048 | -0.117 | 0.021 | 1 | 5552.37 |
| Marine | scaled__year:pollution.habitatl1 | 0.064 | -0.001 | 0.127 | 1 | 3780.68 |
| Marine | scaled__year:pollution.climatechange1 | 0.024 | -0.288 | 0.342 | 1 | 9459.37 |
| Marine | scaled__year:pollution.invasive1 | 0.041 | -0.285 | 0.361 | 1 | 9648.83 |
| Marine | scaled__year:pollution.exploitation1 | 0.072 | 0.012 | 0.130 | 1 | 3646.36 |
| Marine | scaled__year:pollution.disease1 | 0.101 | -0.032 | 0.233 | 1 | 5730.75 |
| Marine | scaled__year:habitatl.climatechange1 | 0.030 | -0.014 | 0.073 | 1 | 4042.20 |
| Marine | scaled__year:habitatl.invasive1 | 0.063 | -0.023 | 0.149 | 1 | 6358.84 |
| Marine | scaled__year:habitatl.exploitation1 | 0.034 | 0.004 | 0.064 | 1 | 4774.11 |
| Marine | scaled__year:habitatl.disease1 | 0.060 | -0.030 | 0.151 | 1 | 5683.81 |
| Marine | scaled__year:climatechange.invasive1 | 0.039 | -0.242 | 0.312 | 1 | 7410.19 |
| Marine | scaled__year:climatechange.exploitation1 | 0.041 | -0.013 | 0.095 | 1 | 5709.23 |
| Marine | scaled__year:invasive.exploitation1 | 0.091 | -0.013 | 0.195 | 1 | 7375.35 |

Table S2: Model coefficents for global population trends across systems. Median represents the median of the posterior distribution. CI low and high are the lower and higher values of the 95% credible interval. Rhat is the ratio of the effective sample size to the overall number of iterations, with values close to one indicating convergence values. *(continued)*

| System | Parameter | Median | CI_low | CI_high | Rhat | ESS |
|---|---|---|---|---|---|---|
| Marine | scaled_year:invasive.disease1 | 0.020 | -0.280 | 0.320 | 1 | 8839.66 |
| Marine | scaled_year:exploitation.disease1 | 0.022 | -0.100 | 0.146 | 1 | 6834.71 |
| Marine | scaled_year:pollution.habitatl.climatechange1 | -0.086 | -0.407 | 0.232 | 1 | 9502.83 |
| Marine | scaled_year:pollution.habitatl.exploitation1 | -0.152 | -0.230 | -0.072 | 1 | 3743.66 |
| Marine | scaled_year:pollution.habitatl.disease1 | -0.115 | -0.286 | 0.052 | 1 | 5532.57 |
| Marine | scaled_year:pollution.climatechange.invasive1 | 0.104 | -0.276 | 0.494 | 1 | 11137.36 |
| Marine | scaled_year:pollution.invasive.exploitation1 | -0.080 | -0.406 | 0.248 | 1 | 9522.38 |
| Marine | scaled_year:pollution.exploitation.disease1 | -0.032 | -0.214 | 0.146 | 1 | 6262.19 |
| Marine | scaled_year:habitatl.climatechange.invasive1 | -0.041 | -0.339 | 0.261 | 1 | 8158.14 |
| Marine | scaled_year:habitatl.climatechange.exploitation1 | -0.054 | -0.134 | 0.028 | 1 | 4861.83 |
| Marine | scaled_year:habitatl.invasive.exploitation1 | -0.031 | -0.178 | 0.122 | 1 | 6852.32 |
| Marine | scaled_year:habitatl.invasive.disease1 | 0.002 | -0.308 | 0.320 | 1 | 9101.91 |
| Marine | scaled_year:habitatl.exploitation.disease1 | -0.011 | -0.205 | 0.184 | 1 | 8130.89 |
| Marine | scaled_year:climatechange.invasive.exploitation1 | -0.035 | -0.328 | 0.255 | 1 | 7910.46 |
| Marine | scaled_year:invasive.exploitation.disease1 | 0.006 | -0.305 | 0.318 | 1 | 9352.43 |
| Terrestrial | Intercept | -0.002 | -0.014 | 0.011 | 1 | 14493.64 |
| Terrestrial | scaled_year | 0.027 | -0.168 | 0.223 | 1 | 4434.99 |
| Terrestrial | pollution1 | 0.005 | -0.101 | 0.109 | 1 | 9374.26 |
| Terrestrial | habitatl1 | 0.002 | -0.029 | 0.032 | 1 | 13425.78 |
| Terrestrial | climatechange1 | 0.001 | -0.070 | 0.071 | 1 | 12328.32 |
| Terrestrial | invasive1 | 0.003 | -0.063 | 0.071 | 1 | 12510.32 |
| Terrestrial | exploitation1 | 0.000 | -0.046 | 0.047 | 1 | 11955.00 |
| Terrestrial | disease1 | 0.005 | -0.077 | 0.087 | 1 | 11480.46 |
| Terrestrial | pollution.habitatl1 | -0.003 | -0.139 | 0.134 | 1 | 9142.36 |
| Terrestrial | pollution.climatechange1 | -0.002 | -0.382 | 0.377 | 1 | 11445.65 |
| Terrestrial | pollution.invasive1 | 0.036 | -0.292 | 0.369 | 1 | 9421.51 |
| Terrestrial | pollution.exploitation1 | -0.022 | -0.161 | 0.118 | 1 | 9204.04 |
| Terrestrial | pollution.disease1 | -0.026 | -0.338 | 0.289 | 1 | 12020.10 |
| Terrestrial | habitatl.climatechange1 | -0.003 | -0.095 | 0.092 | 1 | 11206.66 |
| Terrestrial | habitatl.invasive1 | -0.011 | -0.128 | 0.107 | 1 | 11554.16 |
| Terrestrial | habitatl.exploitation1 | -0.002 | -0.070 | 0.063 | 1 | 10422.58 |
| Terrestrial | habitatl.disease1 | 0.008 | -0.162 | 0.175 | 1 | 11278.90 |
| Terrestrial | climatechange.invasive1 | -0.001 | -0.227 | 0.227 | 1 | 12684.44 |
| Terrestrial | climatechange.exploitation1 | 0.003 | -0.236 | 0.248 | 1 | 11323.32 |
| Terrestrial | climatechange.disease1 | -0.002 | -0.369 | 0.382 | 1 | 12596.79 |
| Terrestrial | invasive.exploitation1 | -0.002 | -0.218 | 0.213 | 1 | 13150.68 |
| Terrestrial | invasive.disease1 | -0.025 | -0.189 | 0.138 | 1 | 12321.57 |
| Terrestrial | exploitation.disease1 | -0.008 | -0.186 | 0.171 | 1 | 12451.48 |
| Terrestrial | pollution.habitatl.climatechange1 | 0.000 | -0.376 | 0.379 | 1 | 10559.86 |
| Terrestrial | pollution.habitatl.invasive1 | -0.022 | -0.371 | 0.337 | 1 | 10092.42 |
| Terrestrial | pollution.habitatl.exploitation1 | 0.022 | -0.238 | 0.282 | 1 | 12526.66 |
| Terrestrial | pollution.habitatl.disease1 | 0.000 | -0.344 | 0.331 | 1 | 11984.15 |
| Terrestrial | pollution.invasive.exploitation1 | 0.047 | -0.345 | 0.432 | 1 | 12009.52 |
| Terrestrial | habitatl.climatechange.invasive1 | 0.031 | -0.251 | 0.310 | 1 | 11881.08 |
| Terrestrial | habitatl.climatechange.exploitation1 | -0.004 | -0.272 | 0.252 | 1 | 10582.68 |
| Terrestrial | habitatl.climatechange.disease1 | -0.002 | -0.386 | 0.363 | 1 | 11457.51 |
| Terrestrial | habitatl.invasive.exploitation1 | -0.008 | -0.283 | 0.278 | 1 | 11930.29 |
| Terrestrial | habitatl.invasive.disease1 | 0.018 | -0.257 | 0.294 | 1 | 11676.28 |
| Terrestrial | habitatl.exploitation.disease1 | -0.010 | -0.269 | 0.249 | 1 | 10326.91 |
| Terrestrial | invasive.exploitation.disease1 | -0.022 | -0.411 | 0.368 | 1 | 17636.14 |
| Terrestrial | scaled_year:pollution1 | -0.024 | -0.079 | 0.029 | 1 | 3268.79 |
| Terrestrial | scaled_year:habitatl1 | -0.028 | -0.043 | -0.014 | 1 | 5017.55 |
| Terrestrial | scaled_year:climatechange1 | -0.069 | -0.103 | -0.037 | 1 | 4880.76 |
| Terrestrial | scaled_year:invasive1 | -0.081 | -0.119 | -0.041 | 1 | 5367.83 |
| Terrestrial | scaled_year:exploitation1 | -0.002 | -0.023 | 0.021 | 1 | 5385.58 |
| Terrestrial | scaled_year:disease1 | -0.043 | -0.083 | -0.002 | 1 | 4499.32 |
| Terrestrial | scaled_year:pollution.habitatl1 | 0.011 | -0.061 | 0.082 | 1 | 3443.67 |

Table S2: Model coefficents for global population trends across systems. Median represents the median of the posterior distribution. CI low and high are the lower and higher values of the 95% credible interval. Rhat is the ratio of the effective sample size to the overall number of iterations, with values close to one indicating convergence values. *(continued)*

| System | Parameter | Median | CI_low | CI_high | Rhat | ESS |
|---|---|---|---|---|---|---|
| Terrestrial | scaled_year:pollution.climatechange1 | -0.001 | -0.361 | 0.354 | 1 | 10361.97 |
| Terrestrial | scaled_year:pollution.invasive1 | 0.005 | -0.288 | 0.300 | 1 | 7397.21 |
| Terrestrial | scaled_year:pollution.exploitation1 | -0.064 | -0.140 | 0.014 | 1 | 3747.35 |
| Terrestrial | scaled_year:pollution.disease1 | -0.242 | -0.408 | -0.071 | 1 | 6403.90 |
| Terrestrial | scaled_year:habitatl.climatechange1 | 0.061 | 0.015 | 0.107 | 1 | 4222.72 |
| Terrestrial | scaled_year:habitatl.invasive1 | 0.058 | -0.003 | 0.119 | 1 | 4876.14 |
| Terrestrial | scaled_year:habitatl.exploitation1 | -0.014 | -0.046 | 0.016 | 1 | 4832.00 |
| Terrestrial | scaled_year:habitatl.disease1 | 0.069 | -0.016 | 0.154 | 1 | 5307.78 |
| Terrestrial | scaled_year:climatechange.invasive1 | 0.090 | -0.059 | 0.238 | 1 | 6733.50 |
| Terrestrial | scaled_year:climatechange.exploitation1 | 0.023 | -0.111 | 0.152 | 1 | 6621.00 |
| Terrestrial | scaled_year:climatechange.disease1 | -0.013 | -0.368 | 0.343 | 1 | 11160.56 |
| Terrestrial | scaled_year:invasive.exploitation1 | -0.042 | -0.156 | 0.071 | 1 | 7719.45 |
| Terrestrial | scaled_year:invasive.disease1 | 0.078 | -0.008 | 0.162 | 1 | 4533.76 |
| Terrestrial | scaled_year:exploitation.disease1 | -0.044 | -0.140 | 0.051 | 1 | 5243.55 |
| Terrestrial | scaled_year:pollution.habitatl.climatechange1 | 0.001 | -0.352 | 0.356 | 1 | 10235.70 |
| Terrestrial | scaled_year:pollution.habitatl.invasive1 | 0.027 | -0.278 | 0.326 | 1 | 7482.68 |
| Terrestrial | scaled_year:pollution.habitatl.exploitation1 | 0.127 | -0.002 | 0.253 | 1 | 4940.92 |
| Terrestrial | scaled_year:pollution.habitatl.disease1 | 0.204 | 0.007 | 0.393 | 1 | 6253.15 |
| Terrestrial | scaled_year:pollution.invasive.exploitation1 | -0.032 | -0.356 | 0.292 | 1 | 7957.89 |
| Terrestrial | scaled_year:habitatl.climatechange.invasive1 | -0.069 | -0.246 | 0.108 | 1 | 6652.07 |
| Terrestrial | scaled_year:habitatl.climatechange.exploitation1 | 0.013 | -0.134 | 0.163 | 1 | 5754.05 |
| Terrestrial | scaled_year:habitatl.climatechange.disease1 | -0.007 | -0.360 | 0.350 | 1 | 11127.80 |
| Terrestrial | scaled_year:habitatl.invasive.exploitation1 | 0.054 | -0.093 | 0.205 | 1 | 7178.80 |
| Terrestrial | scaled_year:habitatl.invasive.disease1 | -0.132 | -0.283 | 0.020 | 1 | 5421.63 |
| Terrestrial | scaled_year:habitatl.exploitation.disease1 | 0.079 | -0.056 | 0.211 | 1 | 5032.60 |
| Terrestrial | scaled_year:invasive.exploitation.disease1 | -0.263 | -0.479 | -0.046 | 1 | 7547.80 |

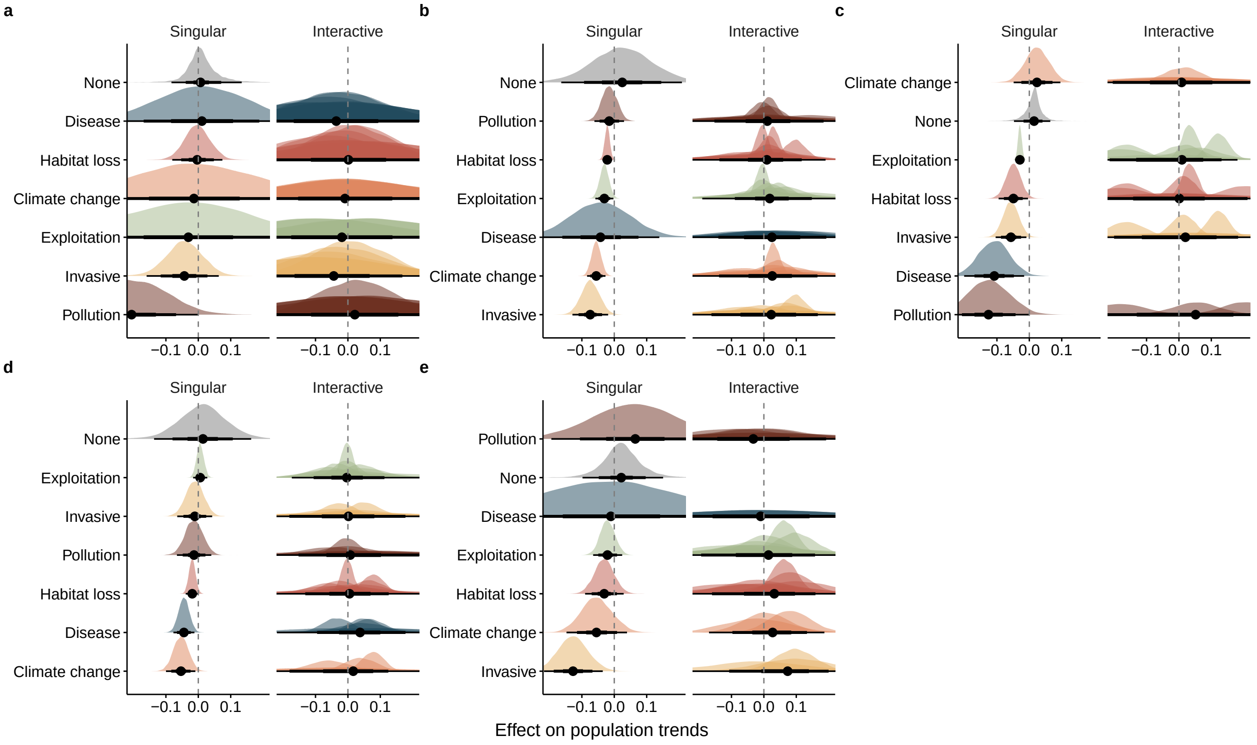


Figure S4: Effects of threats on vertebrate population trends across taxa. 95 % credible intervals of the effects of single and interacting threats upon (a) amphibian, (b) bird, (c) fish, (d) mammalian, and (e) reptilian time series trends. The dashed vertical line shows zero influence – i.e., no effect of the factors – while the None parameter is the trend in the absence of threats. The remaining parameters represent modifications of this None trend. The credible intervals are based on 1,000 samples from the posterior distribution of the model coefficients

Table S3: Model coefficents for global population trends across taxa. Median represents the median of the posterior distribution. CI low and high are the lower and higher values of the 95% credible interval. Rhat is the ratio of the effective sample size to the overall number of iterations, with values close to one indicating convergence values.

| Taxon | Parameter | Median | CI_low | CI_high | Rhat | ESS |
|---|---|---|---|---|---|---|
| Amphibian | Intercept | -0.001 | -0.066 | 0.064 | 1 | 14579.36 |
| Amphibian | scaled_year | 0.007 | -0.083 | 0.134 | 1 | 3694.44 |
| Amphibian | pollution1 | -0.005 | -0.300 | 0.289 | 1 | 9882.46 |
| Amphibian | habitatl1 | -0.002 | -0.137 | 0.136 | 1 | 13599.16 |
| Amphibian | climatechange1 | 0.002 | -0.410 | 0.425 | 1 | 12117.86 |
| Amphibian | invasive1 | -0.008 | -0.172 | 0.156 | 1 | 14231.75 |
| Amphibian | exploitation1 | 0.007 | -0.401 | 0.410 | 1 | 13772.79 |
| Amphibian | disease1 | 0.007 | -0.304 | 0.325 | 1 | 8199.11 |
| Amphibian | pollution.habitatl1 | -0.018 | -0.346 | 0.309 | 1 | 9290.12 |
| Amphibian | pollution.climatechange1 | 0.003 | -0.440 | 0.456 | 1 | 11816.81 |
| Amphibian | pollution.exploitation1 | 0.012 | -0.440 | 0.448 | 1 | 14004.20 |
| Amphibian | pollution.disease1 | 0.003 | -0.430 | 0.449 | 1 | 12466.40 |
| Amphibian | habitatl.climatechange1 | -0.005 | -0.427 | 0.421 | 1 | 13770.74 |
| Amphibian | habitatl.invasive1 | 0.008 | -0.330 | 0.347 | 1 | 12400.51 |
| Amphibian | habitatl.exploitation1 | 0.009 | -0.440 | 0.454 | 1 | 12170.11 |
| Amphibian | habitatl.disease1 | 0.007 | -0.314 | 0.337 | 1 | 8708.66 |
| Amphibian | climatechange.disease1 | 0.004 | -0.418 | 0.414 | 1 | 13073.73 |
| Amphibian | invasive.exploitation1 | -0.012 | -0.464 | 0.436 | 1 | 14937.05 |
| Amphibian | invasive.disease1 | -0.017 | -0.356 | 0.321 | 1 | 10051.85 |
| Amphibian | exploitation.disease1 | -0.016 | -0.467 | 0.447 | 1 | 15948.48 |
| Amphibian | pollution.habitatl.exploitation1 | 0.012 | -0.429 | 0.449 | 1 | 14701.47 |
| Amphibian | pollution.climatechange.disease1 | 0.005 | -0.449 | 0.466 | 1 | 13396.04 |
| Amphibian | habitatl.climatechange.disease1 | -0.008 | -0.432 | 0.417 | 1 | 12256.46 |
| Amphibian | habitatl.invasive.disease1 | 0.007 | -0.400 | 0.403 | 1 | 13050.00 |
| Amphibian | invasive.exploitation.disease1 | -0.013 | -0.460 | 0.434 | 1 | 15033.52 |
| Amphibian | scaled_year:pollution1 | -0.206 | -0.418 | 0.000 | 1 | 6658.07 |
| Amphibian | scaled_year:habitatl1 | -0.004 | -0.080 | 0.074 | 1 | 4933.38 |
| Amphibian | scaled_year:climatechange1 | -0.013 | -0.422 | 0.393 | 1 | 12429.84 |
| Amphibian | scaled_year:invasive1 | -0.043 | -0.159 | 0.063 | 1 | 4073.63 |
| Amphibian | scaled_year:exploitation1 | -0.031 | -0.428 | 0.363 | 1 | 11943.78 |
| Amphibian | scaled_year:disease1 | 0.011 | -0.266 | 0.282 | 1 | 5081.27 |
| Amphibian | scaled_year:pollution.habitatl1 | 0.027 | -0.201 | 0.257 | 1 | 6207.57 |
| Amphibian | scaled_year:pollution.climatechange1 | -0.004 | -0.452 | 0.425 | 1 | 11973.56 |
| Amphibian | scaled_year:pollution.exploitation1 | 0.055 | -0.372 | 0.486 | 1 | 12214.14 |
| Amphibian | scaled_year:pollution.disease1 | -0.006 | -0.442 | 0.438 | 1 | 13487.54 |
| Amphibian | scaled_year:habitatl.climatechange1 | -0.016 | -0.425 | 0.400 | 1 | 12998.10 |
| Amphibian | scaled_year:habitatl.invasive1 | 0.007 | -0.222 | 0.238 | 1 | 6750.96 |
| Amphibian | scaled_year:habitatl.exploitation1 | 0.056 | -0.362 | 0.487 | 1 | 13253.52 |
| Amphibian | scaled_year:habitatl.disease1 | -0.027 | -0.311 | 0.249 | 1 | 5498.01 |
| Amphibian | scaled_year:climatechange.disease1 | -0.008 | -0.423 | 0.408 | 1 | 11342.39 |
| Amphibian | scaled_year:invasive.exploitation1 | -0.091 | -0.523 | 0.340 | 1 | 12842.61 |
| Amphibian | scaled_year:invasive.disease1 | -0.026 | -0.298 | 0.250 | 1 | 5245.63 |
| Amphibian | scaled_year:exploitation.disease1 | -0.094 | -0.519 | 0.339 | 1 | 12950.98 |
| Amphibian | scaled_year:pollution.habitatl.exploitation1 | 0.054 | -0.378 | 0.492 | 1 | 13537.20 |
| Amphibian | scaled_year:pollution.climatechange.disease1 | -0.006 | -0.433 | 0.427 | 1 | 12123.34 |
| Amphibian | scaled_year:habitatl.climatechange.disease1 | -0.011 | -0.424 | 0.406 | 1 | 10203.33 |
| Amphibian | scaled_year:habitatl.invasive.disease1 | -0.065 | -0.398 | 0.278 | 1 | 7533.38 |
| Amphibian | scaled_year:invasive.exploitation.disease1 | -0.092 | -0.523 | 0.349 | 1 | 13267.58 |
| Bird | Intercept | -0.002 | -0.016 | 0.011 | 1 | 16717.18 |
| Bird | scaled_year | 0.025 | -0.164 | 0.209 | 1 | 10510.46 |
| Bird | pollution1 | -0.001 | -0.081 | 0.078 | 1 | 12422.21 |
| Bird | habitatl1 | -0.002 | -0.029 | 0.025 | 1 | 19065.69 |
| Bird | climatechange1 | 0.000 | -0.054 | 0.053 | 1 | 19004.58 |
| Bird | invasive1 | 0.010 | -0.063 | 0.082 | 1 | 17879.56 |
| Bird | exploitation1 | 0.002 | -0.055 | 0.055 | 1 | 13831.46 |
| Bird | disease1 | -0.005 | -0.213 | 0.207 | 1 | 8187.61 |
| Bird | pollution.habitatl1 | 0.006 | -0.088 | 0.100 | 1 | 12197.52 |

Table S3: Model coefficents for global population trends across taxa. Median represents the median of the posterior distribution. CI low and high are the lower and higher values of the 95% credible interval. Rhat is the ratio of the effective sample size to the overall number of iterations, with values close to one indicating convergence values. *(continued)*

| Taxon | Parameter | Median | CI_low | CI_high | Rhat | ESS |
|---|---|---|---|---|---|---|
| Bird | pollution.climatechange1 | -0.003 | -0.318 | 0.317 | 1 | 16348.85 |
| Bird | pollution.invasive1 | 0.012 | -0.264 | 0.293 | 1 | 14153.07 |
| Bird | pollution.exploitation1 | -0.013 | -0.125 | 0.104 | 1 | 10420.23 |
| Bird | pollution.disease1 | -0.001 | -0.239 | 0.235 | 1 | 9526.25 |
| Bird | habitatl.climatechange1 | 0.001 | -0.077 | 0.078 | 1 | 17127.92 |
| Bird | habitatl.invasive1 | -0.007 | -0.113 | 0.098 | 1 | 16592.59 |
| Bird | habitatl.exploitation1 | 0.003 | -0.071 | 0.080 | 1 | 14004.14 |
| Bird | habitatl.disease1 | 0.011 | -0.195 | 0.224 | 1 | 8619.98 |
| Bird | climatechange.invasive1 | -0.009 | -0.225 | 0.206 | 1 | 16500.64 |
| Bird | climatechange.exploitation1 | 0.003 | -0.149 | 0.158 | 1 | 17521.60 |
| Bird | climatechange.disease1 | -0.006 | -0.258 | 0.247 | 1 | 11131.63 |
| Bird | invasive.exploitation1 | 0.002 | -0.183 | 0.190 | 1 | 16556.68 |
| Bird | invasive.disease1 | 0.000 | -0.317 | 0.316 | 1 | 15972.68 |
| Bird | exploitation.disease1 | 0.000 | -0.249 | 0.255 | 1 | 12401.71 |
| Bird | pollution.habitatl.climatechange1 | 0.001 | -0.333 | 0.324 | 1 | 15654.29 |
| Bird | pollution.habitatl.invasive1 | -0.008 | -0.315 | 0.291 | 1 | 14886.59 |
| Bird | pollution.habitatl.exploitation1 | 0.001 | -0.149 | 0.150 | 1 | 10358.20 |
| Bird | pollution.habitatl.disease1 | -0.013 | -0.269 | 0.240 | 1 | 10514.22 |
| Bird | pollution.climatechange.invasive1 | -0.008 | -0.395 | 0.387 | 1 | 15438.46 |
| Bird | pollution.invasive.exploitation1 | 0.015 | -0.310 | 0.332 | 1 | 13564.28 |
| Bird | pollution.exploitation.disease1 | 0.024 | -0.280 | 0.331 | 1 | 14111.12 |
| Bird | habitatl.climatechange.invasive1 | 0.012 | -0.262 | 0.282 | 1 | 16145.15 |
| Bird | habitatl.climatechange.exploitation1 | -0.005 | -0.215 | 0.208 | 1 | 17138.97 |
| Bird | habitatl.climatechange.disease1 | 0.001 | -0.285 | 0.290 | 1 | 13146.66 |
| Bird | habitatl.invasive.exploitation1 | 0.000 | -0.237 | 0.232 | 1 | 15914.43 |
| Bird | habitatl.invasive.disease1 | -0.005 | -0.341 | 0.329 | 1 | 16488.54 |
| Bird | habitatl.exploitation.disease1 | -0.008 | -0.311 | 0.291 | 1 | 14134.10 |
| Bird | climatechange.invasive.exploitation1 | -0.002 | -0.336 | 0.321 | 1 | 18670.21 |
| Bird | invasive.exploitation.disease1 | -0.003 | -0.382 | 0.366 | 1 | 16966.73 |
| Bird | scaled_year:pollution1 | -0.016 | -0.061 | 0.030 | 1 | 5814.25 |
| Bird | scaled_year:habitatl1 | -0.022 | -0.037 | -0.007 | 1 | 6850.22 |
| Bird | scaled_year:climatechange1 | -0.056 | -0.084 | -0.027 | 1 | 6772.98 |
| Bird | scaled_year:invasive1 | -0.074 | -0.129 | -0.019 | 1 | 6534.58 |
| Bird | scaled_year:exploitation1 | -0.031 | -0.058 | -0.003 | 1 | 6741.07 |
| Bird | scaled_year:disease1 | -0.043 | -0.225 | 0.139 | 1 | 5267.55 |
| Bird | scaled_year:pollution.habitatl1 | 0.015 | -0.040 | 0.069 | 1 | 5686.33 |
| Bird | scaled_year:pollution.climatechange1 | 0.051 | -0.261 | 0.350 | 1 | 14565.93 |
| Bird | scaled_year:pollution.invasive1 | 0.003 | -0.269 | 0.278 | 1 | 10709.03 |
| Bird | scaled_year:pollution.exploitation1 | -0.006 | -0.073 | 0.063 | 1 | 5918.79 |
| Bird | scaled_year:pollution.disease1 | -0.015 | -0.213 | 0.172 | 1 | 5584.58 |
| Bird | scaled_year:habitatl.climatechange1 | 0.029 | -0.010 | 0.069 | 1 | 6413.97 |
| Bird | scaled_year:habitatl.invasive1 | 0.096 | 0.029 | 0.163 | 1 | 6866.55 |
| Bird | scaled_year:habitatl.exploitation1 | -0.005 | -0.043 | 0.033 | 1 | 6490.84 |
| Bird | scaled_year:habitatl.disease1 | 0.047 | -0.133 | 0.235 | 1 | 5401.86 |
| Bird | scaled_year:climatechange.invasive1 | 0.072 | -0.073 | 0.216 | 1 | 10272.22 |
| Bird | scaled_year:climatechange.exploitation1 | 0.043 | -0.040 | 0.125 | 1 | 8899.76 |
| Bird | scaled_year:climatechange.disease1 | 0.091 | -0.114 | 0.292 | 1 | 6247.30 |
| Bird | scaled_year:invasive.exploitation1 | 0.068 | -0.041 | 0.180 | 1 | 9095.51 |
| Bird | scaled_year:invasive.disease1 | 0.016 | -0.281 | 0.316 | 1 | 13995.67 |
| Bird | scaled_year:exploitation.disease1 | -0.012 | -0.205 | 0.185 | 1 | 5865.03 |
| Bird | scaled_year:pollution.habitatl.climatechange1 | -0.053 | -0.362 | 0.263 | 1 | 14646.26 |
| Bird | scaled_year:pollution.habitatl.invasive1 | -0.069 | -0.351 | 0.210 | 1 | 10682.42 |
| Bird | scaled_year:pollution.habitatl.exploitation1 | 0.015 | -0.069 | 0.100 | 1 | 6068.10 |
| Bird | scaled_year:pollution.habitatl.disease1 | 0.012 | -0.192 | 0.211 | 1 | 5968.07 |
| Bird | scaled_year:pollution.climatechange.invasive1 | 0.098 | -0.267 | 0.453 | 1 | 14805.78 |
| Bird | scaled_year:pollution.invasive.exploitation1 | -0.033 | -0.322 | 0.256 | 1 | 11241.73 |
| Bird | scaled_year:pollution.exploitation.disease1 | 0.145 | -0.077 | 0.370 | 1 | 7067.80 |
| Bird | scaled_year:habitatl.climatechange.invasive1 | -0.065 | -0.236 | 0.108 | 1 | 10899.09 |

Table S3: Model coefficents for global population trends across taxa. Median represents the median of the posterior distribution. CI low and high are the lower and higher values of the 95% credible interval. Rhat is the ratio of the effective sample size to the overall number of iterations, with values close to one indicating convergence values. *(continued)*

| Taxon | Parameter | Median | CI_low | CI_high | Rhat | ESS |
|---|---|---|---|---|---|---|
| Bird | scaled_year:habitatl.climatechange.exploitation1 | 0.033 | -0.083 | 0.154 | 1 | 9777.83 |
| Bird | scaled_year:habitatl.climatechange.disease1 | -0.060 | -0.275 | 0.159 | 1 | 7040.76 |
| Bird | scaled_year:habitatl.invasive.exploitation1 | -0.009 | -0.149 | 0.128 | 1 | 8652.08 |
| Bird | scaled_year:habitatl.invasive.disease1 | -0.044 | -0.349 | 0.263 | 1 | 13905.08 |
| Bird | scaled_year:habitatl.exploitation.disease1 | 0.041 | -0.173 | 0.260 | 1 | 6775.46 |
| Bird | scaled_year:climatechange.invasive.exploitation1 | -0.015 | -0.234 | 0.207 | 1 | 11913.75 |
| Bird | scaled_year:invasive.exploitation.disease1 | 0.052 | -0.260 | 0.369 | 1 | 14335.59 |
| Fish | Intercept | 0.006 | -0.018 | 0.030 | 1 | 16683.51 |
| Fish | scaled_year | 0.015 | -0.047 | 0.065 | 1 | 3715.55 |
| Fish | pollution1 | -0.014 | -0.229 | 0.206 | 1 | 6291.72 |
| Fish | habitatl1 | -0.046 | -0.163 | 0.069 | 1 | 7616.80 |
| Fish | climatechange1 | -0.086 | -0.252 | 0.088 | 1 | 12008.58 |
| Fish | invasive1 | -0.025 | -0.144 | 0.091 | 1 | 8158.37 |
| Fish | exploitation1 | -0.012 | -0.042 | 0.018 | 1 | 16879.51 |
| Fish | disease1 | -0.051 | -0.275 | 0.172 | 1 | 17495.29 |
| Fish | pollution.habitatl1 | 0.052 | -0.188 | 0.286 | 1 | 6188.59 |
| Fish | pollution.exploitation1 | -0.067 | -0.311 | 0.167 | 1 | 6377.71 |
| Fish | habitatl.climatechange1 | 0.015 | -0.318 | 0.357 | 1 | 9189.12 |
| Fish | habitatl.invasive1 | 0.069 | -0.099 | 0.239 | 1 | 6926.90 |
| Fish | habitatl.exploitation1 | 0.036 | -0.093 | 0.165 | 1 | 7599.65 |
| Fish | climatechange.invasive1 | 0.008 | -0.388 | 0.408 | 1 | 11575.30 |
| Fish | climatechange.exploitation1 | 0.060 | -0.199 | 0.319 | 1 | 12378.64 |
| Fish | invasive.exploitation1 | 0.018 | -0.135 | 0.168 | 1 | 8219.79 |
| Fish | pollution.habitatl.exploitation1 | 0.036 | -0.252 | 0.329 | 1 | 7468.88 |
| Fish | habitatl.climatechange.invasive1 | 0.008 | -0.395 | 0.405 | 1 | 12068.20 |
| Fish | habitatl.climatechange.exploitation1 | -0.005 | -0.360 | 0.340 | 1 | 8969.69 |
| Fish | habitatl.invasive.exploitation1 | -0.041 | -0.295 | 0.221 | 1 | 9326.49 |
| Fish | scaled_year:pollution1 | -0.126 | -0.251 | -0.001 | 1 | 4771.62 |
| Fish | scaled_year:habitatl1 | -0.049 | -0.095 | -0.004 | 1 | 4269.67 |
| Fish | scaled_year:climatechange1 | 0.023 | -0.048 | 0.096 | 1 | 7835.41 |
| Fish | scaled_year:invasive1 | -0.057 | -0.104 | -0.009 | 1 | 4430.12 |
| Fish | scaled_year:exploitation1 | -0.029 | -0.041 | -0.018 | 1 | 3048.03 |
| Fish | scaled_year:disease1 | -0.108 | -0.201 | -0.016 | 1 | 7999.89 |
| Fish | scaled_year:pollution.habitatl1 | 0.203 | 0.062 | 0.340 | 1 | 4508.85 |
| Fish | scaled_year:pollution.exploitation1 | 0.054 | -0.086 | 0.200 | 1 | 4458.06 |
| Fish | scaled_year:habitatl.climatechange1 | -0.004 | -0.324 | 0.318 | 1 | 8167.18 |
| Fish | scaled_year:habitatl.invasive1 | 0.014 | -0.063 | 0.093 | 1 | 3860.06 |
| Fish | scaled_year:habitatl.exploitation1 | 0.034 | -0.015 | 0.084 | 1 | 4118.54 |
| Fish | scaled_year:climatechange.invasive1 | 0.037 | -0.359 | 0.422 | 1 | 10804.90 |
| Fish | scaled_year:climatechange.exploitation1 | 0.014 | -0.103 | 0.132 | 1 | 6748.65 |
| Fish | scaled_year:invasive.exploitation1 | 0.121 | 0.055 | 0.188 | 1 | 4282.73 |
| Fish | scaled_year:pollution.habitatl.exploitation1 | -0.184 | -0.343 | -0.023 | 1 | 4507.96 |
| Fish | scaled_year:habitatl.climatechange.invasive1 | 0.036 | -0.354 | 0.429 | 1 | 11669.59 |
| Fish | scaled_year:habitatl.climatechange.exploitation1 | -0.039 | -0.369 | 0.286 | 1 | 8267.37 |
| Fish | scaled_year:habitatl.invasive.exploitation1 | -0.163 | -0.280 | -0.054 | 1 | 4456.38 |
| Mammal | Intercept | -0.003 | -0.021 | 0.016 | 1 | 13849.69 |
| Mammal | scaled_year | 0.014 | -0.136 | 0.163 | 1 | 6335.64 |
| Mammal | pollution1 | 0.008 | -0.134 | 0.154 | 1 | 12538.67 |
| Mammal | habitatl1 | 0.005 | -0.046 | 0.055 | 1 | 12550.43 |
| Mammal | climatechange1 | 0.004 | -0.113 | 0.122 | 1 | 13397.77 |
| Mammal | invasive1 | 0.006 | -0.109 | 0.121 | 1 | 13585.03 |
| Mammal | exploitation1 | -0.001 | -0.056 | 0.055 | 1 | 12603.97 |
| Mammal | disease1 | 0.006 | -0.079 | 0.092 | 1 | 15148.06 |
| Mammal | pollution.habitatl1 | -0.001 | -0.276 | 0.272 | 1 | 12096.11 |
| Mammal | pollution.invasive1 | -0.006 | -0.423 | 0.403 | 1 | 15459.25 |
| Mammal | pollution.exploitation1 | -0.008 | -0.216 | 0.195 | 1 | 12696.07 |
| Mammal | pollution.disease1 | -0.005 | -0.337 | 0.330 | 1 | 11905.51 |

Table S3: Model coefficents for global population trends across taxa. Median represents the median of the posterior distribution. CI low and high are the lower and higher values of the 95% credible interval. Rhat is the ratio of the effective sample size to the overall number of iterations, with values close to one indicating convergence values. *(continued)*

| Taxon | Parameter | Median | CI_low | CI_high | Rhat | ESS |
|---|---|---|---|---|---|---|
| Mammal | habitatl.climatechange1 | 0.006 | -0.163 | 0.172 | 1 | 12451.62 |
| Mammal | habitatl.invasive1 | -0.012 | -0.219 | 0.202 | 1 | 14760.28 |
| Mammal | habitatl.exploitation1 | -0.001 | -0.098 | 0.097 | 1 | 11439.19 |
| Mammal | habitatl.disease1 | -0.008 | -0.244 | 0.226 | 1 | 14406.44 |
| Mammal | climatechange.invasive1 | 0.043 | -0.374 | 0.455 | 1 | 15330.09 |
| Mammal | climatechange.exploitation1 | -0.005 | -0.236 | 0.229 | 1 | 12960.58 |
| Mammal | invasive.exploitation1 | -0.006 | -0.325 | 0.310 | 1 | 16644.23 |
| Mammal | invasive.disease1 | -0.029 | -0.247 | 0.180 | 1 | 12164.63 |
| Mammal | exploitation.disease1 | 0.001 | -0.207 | 0.199 | 1 | 14913.80 |
| Mammal | pollution.habitatl.invasive1 | -0.005 | -0.396 | 0.403 | 1 | 14763.66 |
| Mammal | pollution.habitatl.exploitation1 | 0.004 | -0.304 | 0.316 | 1 | 11095.23 |
| Mammal | pollution.habitatl.disease1 | -0.011 | -0.399 | 0.379 | 1 | 12632.84 |
| Mammal | pollution.exploitation.disease1 | -0.009 | -0.379 | 0.383 | 1 | 13231.85 |
| Mammal | habitatl.climatechange.invasive1 | 0.038 | -0.377 | 0.449 | 1 | 15400.49 |
| Mammal | habitatl.climatechange.exploitation1 | -0.011 | -0.282 | 0.272 | 1 | 11722.41 |
| Mammal | habitatl.invasive.exploitation1 | -0.011 | -0.359 | 0.338 | 1 | 14750.06 |
| Mammal | habitatl.exploitation.disease1 | -0.010 | -0.316 | 0.294 | 1 | 11029.71 |
| Mammal | scaled_year:pollution1 | -0.013 | -0.066 | 0.040 | 1 | 8083.43 |
| Mammal | scaled_year:habitatl1 | -0.019 | -0.039 | 0.001 | 1 | 7068.25 |
| Mammal | scaled_year:climatechange1 | -0.054 | -0.100 | -0.009 | 1 | 6433.52 |
| Mammal | scaled_year:invasive1 | -0.011 | -0.065 | 0.043 | 1 | 7280.79 |
| Mammal | scaled_year:exploitation1 | 0.006 | -0.016 | 0.028 | 1 | 7216.97 |
| Mammal | scaled_year:disease1 | -0.044 | -0.077 | -0.012 | 1 | 6331.93 |
| Mammal | scaled_year:pollution.habitatl1 | -0.001 | -0.146 | 0.143 | 1 | 8285.16 |
| Mammal | scaled_year:pollution.invasive1 | 0.043 | -0.330 | 0.403 | 1 | 12613.31 |
| Mammal | scaled_year:pollution.exploitation1 | -0.005 | -0.082 | 0.074 | 1 | 8791.91 |
| Mammal | scaled_year:pollution.disease1 | 0.069 | -0.227 | 0.364 | 1 | 8004.59 |
| Mammal | scaled_year:habitatl.climatechange1 | 0.078 | 0.009 | 0.144 | 1 | 6868.76 |
| Mammal | scaled_year:habitatl.invasive1 | -0.027 | -0.124 | 0.070 | 1 | 7884.33 |
| Mammal | scaled_year:habitatl.exploitation1 | -0.003 | -0.041 | 0.035 | 1 | 5385.86 |
| Mammal | scaled_year:habitatl.disease1 | 0.064 | -0.032 | 0.160 | 1 | 8339.05 |
| Mammal | scaled_year:climatechange.invasive1 | -0.040 | -0.406 | 0.317 | 1 | 11810.95 |
| Mammal | scaled_year:climatechange.exploitation1 | 0.038 | -0.060 | 0.138 | 1 | 6350.40 |
| Mammal | scaled_year:invasive.exploitation1 | -0.042 | -0.216 | 0.134 | 1 | 10452.96 |
| Mammal | scaled_year:invasive.disease1 | 0.050 | -0.038 | 0.144 | 1 | 6312.30 |
| Mammal | scaled_year:exploitation.disease1 | -0.033 | -0.121 | 0.058 | 1 | 7024.04 |
| Mammal | scaled_year:pollution.habitatl.invasive1 | 0.044 | -0.322 | 0.412 | 1 | 12659.47 |
| Mammal | scaled_year:pollution.habitatl.exploitation1 | -0.052 | -0.216 | 0.111 | 1 | 8572.24 |
| Mammal | scaled_year:pollution.habitatl.disease1 | -0.040 | -0.347 | 0.277 | 1 | 8442.23 |
| Mammal | scaled_year:pollution.exploitation.disease1 | 0.109 | -0.205 | 0.415 | 1 | 8176.10 |
| Mammal | scaled_year:habitatl.climatechange.invasive1 | -0.043 | -0.407 | 0.317 | 1 | 11424.83 |
| Mammal | scaled_year:habitatl.climatechange.exploitation1 | -0.059 | -0.184 | 0.066 | 1 | 5472.69 |
| Mammal | scaled_year:habitatl.invasive.exploitation1 | 0.023 | -0.179 | 0.230 | 1 | 9403.68 |
| Mammal | scaled_year:habitatl.exploitation.disease1 | 0.046 | -0.090 | 0.184 | 1 | 6512.25 |
| Reptile | Intercept | -0.002 | -0.045 | 0.043 | 1 | 9403.21 |
| Reptile | scaled_year | 0.022 | -0.098 | 0.151 | 1 | 6275.05 |
| Reptile | pollution1 | -0.001 | -0.258 | 0.247 | 1 | 6432.44 |
| Reptile | habitatl1 | 0.003 | -0.058 | 0.064 | 1 | 9122.16 |
| Reptile | climatechange1 | -0.002 | -0.123 | 0.122 | 1 | 10525.23 |
| Reptile | invasive1 | 0.000 | -0.121 | 0.122 | 1 | 14371.84 |
| Reptile | exploitation1 | 0.001 | -0.057 | 0.057 | 1 | 10087.68 |
| Reptile | disease1 | -0.002 | -0.430 | 0.428 | 1 | 23204.65 |
| Reptile | pollution.habitatl1 | 0.004 | -0.251 | 0.254 | 1 | 6535.76 |
| Reptile | pollution.exploitation1 | 0.001 | -0.249 | 0.255 | 1 | 6399.70 |
| Reptile | pollution.disease1 | -0.002 | -0.440 | 0.447 | 1 | 19390.75 |
| Reptile | habitatl.climatechange1 | 0.000 | -0.153 | 0.152 | 1 | 10583.47 |
| Reptile | habitatl.invasive1 | -0.005 | -0.296 | 0.288 | 1 | 17859.94 |
| Reptile | habitatl.exploitation1 | -0.004 | -0.096 | 0.086 | 1 | 8462.92 |

Table S3: Model coefficents for global population trends across taxa. Median represents the median of the posterior distribution. CI low and high are the lower and higher values of the 95% credible interval. Rhat is the ratio of the effective sample size to the overall number of iterations, with values close to one indicating convergence values. *(continued)*

| Taxon | Parameter | Median | CI_low | CI_high | Rhat | ESS |
|---|---|---|---|---|---|---|
| Reptile | climatechange.exploitation1 | 0.005 | -0.186 | 0.196 | 1 | 11827.55 |
| Reptile | invasive.exploitation1 | -0.003 | -0.172 | 0.170 | 1 | 13565.71 |
| Reptile | exploitation.disease1 | 0.000 | -0.439 | 0.426 | 1 | 19662.54 |
| Reptile | pollution.habitatl.exploitation1 | 0.000 | -0.263 | 0.271 | 1 | 6823.35 |
| Reptile | pollution.exploitation.disease1 | 0.000 | -0.441 | 0.445 | 1 | 20507.86 |
| Reptile | habitatl.climatechange.exploitation1 | -0.001 | -0.256 | 0.254 | 1 | 12761.64 |
| Reptile | habitatl.invasive.exploitation1 | 0.000 | -0.350 | 0.350 | 1 | 16846.98 |
| Reptile | scaled_year:pollution1 | 0.065 | -0.194 | 0.321 | 1 | 6678.58 |
| Reptile | scaled_year:habitatl1 | -0.031 | -0.090 | 0.029 | 1 | 4620.64 |
| Reptile | scaled_year:climatechange1 | -0.055 | -0.147 | 0.039 | 1 | 6422.67 |
| Reptile | scaled_year:invasive1 | -0.127 | -0.217 | -0.036 | 1 | 4336.09 |
| Reptile | scaled_year:exploitation1 | -0.021 | -0.065 | 0.024 | 1 | 5205.74 |
| Reptile | scaled_year:disease1 | -0.011 | -0.437 | 0.424 | 1 | 21747.24 |
| Reptile | scaled_year:pollution.habitatl1 | -0.003 | -0.262 | 0.252 | 1 | 7018.00 |
| Reptile | scaled_year:pollution.exploitation1 | -0.027 | -0.286 | 0.232 | 1 | 6692.22 |
| Reptile | scaled_year:pollution.disease1 | -0.008 | -0.441 | 0.412 | 1 | 20641.41 |
| Reptile | scaled_year:habitatl.climatechange1 | 0.080 | -0.043 | 0.206 | 1 | 5429.01 |
| Reptile | scaled_year:habitatl.invasive1 | 0.099 | -0.122 | 0.319 | 1 | 9673.54 |
| Reptile | scaled_year:habitatl.exploitation1 | 0.062 | -0.017 | 0.141 | 1 | 5023.91 |
| Reptile | scaled_year:climatechange.exploitation1 | 0.006 | -0.130 | 0.144 | 1 | 6752.94 |
| Reptile | scaled_year:invasive.exploitation1 | 0.096 | -0.030 | 0.220 | 1 | 7399.39 |
| Reptile | scaled_year:exploitation.disease1 | -0.010 | -0.435 | 0.419 | 1 | 20215.68 |
| Reptile | scaled_year:pollution.habitatl.exploitation1 | -0.095 | -0.359 | 0.167 | 1 | 7013.51 |
| Reptile | scaled_year:pollution.exploitation.disease1 | -0.013 | -0.436 | 0.413 | 1 | 18974.09 |
| Reptile | scaled_year:habitatl.climatechange.exploitation1 | -0.022 | -0.215 | 0.173 | 1 | 6982.58 |
| Reptile | scaled_year:habitatl.invasive.exploitation1 | -0.019 | -0.299 | 0.266 | 1 | 10276.18 |

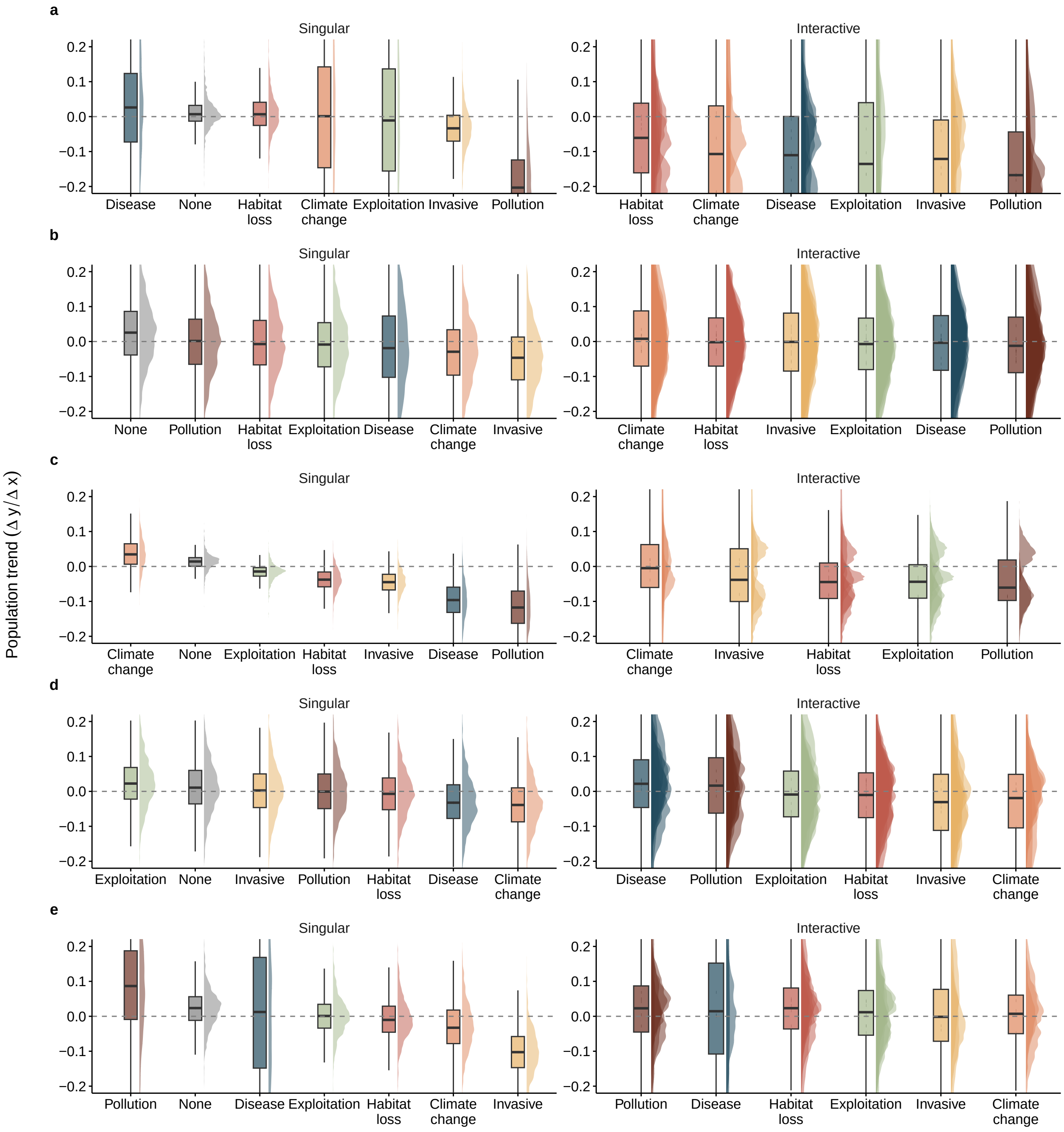


Figure S5: Ultimate effect of threats on vertebrate population trends across taxa. 95 % credible intervals of ultimate estimated trends in the presence of single and interacting threats for (a) amphibian, (b) bird, (c) fish, (d) mammalian, and (e) reptilian. The dashed horizontal line shows the zero-slope – i.e., no effect of the factors. The credible intervals are based on 1,000 samples from the posterior distribution of the model predictions where threats are included/excluded from the model matrix.

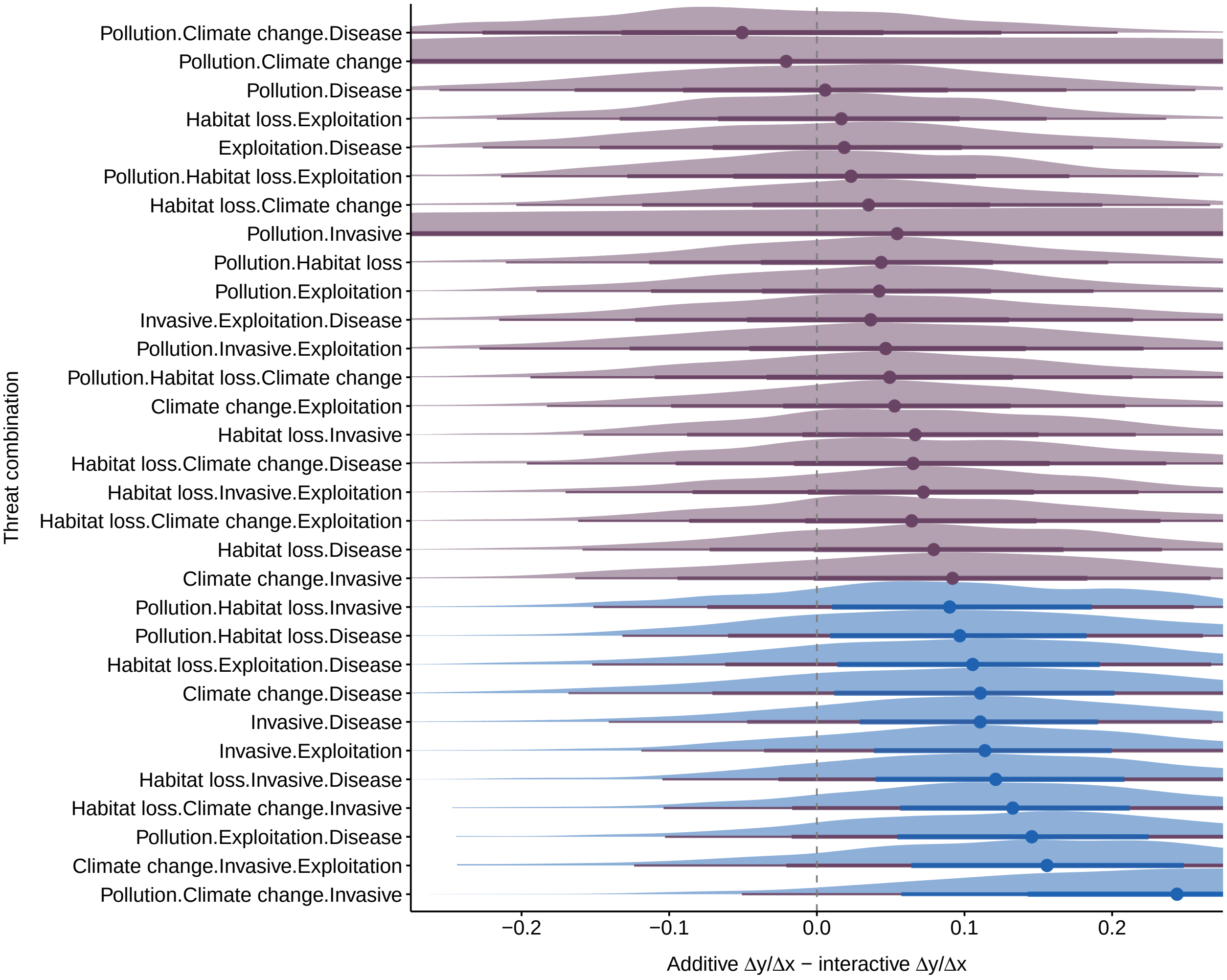


Figure S6: Definition of interacting threat classes using 1000 posterior draws estimated from additive vs interacting model matrices. The difference between additive and interacting trend posteriors is calculated with zero differences assumed to be additive. If the 80 % credible interval does not intersect 0, and is negative, then that threat combination is assumed to be synergistic, whereas if positive, the combination is classified as antagonistic.

Table S4: Proportion of threat interaction types estimated by the global model. n represents the number of interaction types present. Frequency represents the proportion of that given interaction type for that threat and system.

| Threat | Interaction type | n | Frequency |
|---|---|---|---|
| Climate change | Synergistic | 0 | 0.000 |
| Climate change | Antagonistic | 14 | 0.151 |
| Climate change | Additive | 79 | 0.849 |
| Disease | Synergistic | 0 | 0.000 |
| Disease | Antagonistic | 16 | 0.172 |
| Disease | Additive | 77 | 0.828 |
| Exploitation | Synergistic | 0 | 0.000 |
| Exploitation | Antagonistic | 11 | 0.108 |
| Exploitation | Additive | 91 | 0.892 |
| Habitat loss | Synergistic | 0 | 0.000 |
| Habitat loss | Antagonistic | 15 | 0.125 |
| Habitat loss | Additive | 105 | 0.875 |
| Invasive | Synergistic | 0 | 0.000 |
| Invasive | Antagonistic | 22 | 0.216 |
| Invasive | Additive | 80 | 0.784 |
| Pollution | Synergistic | 0 | 0.000 |
| Pollution | Antagonistic | 15 | 0.147 |
| Pollution | Additive | 87 | 0.853 |

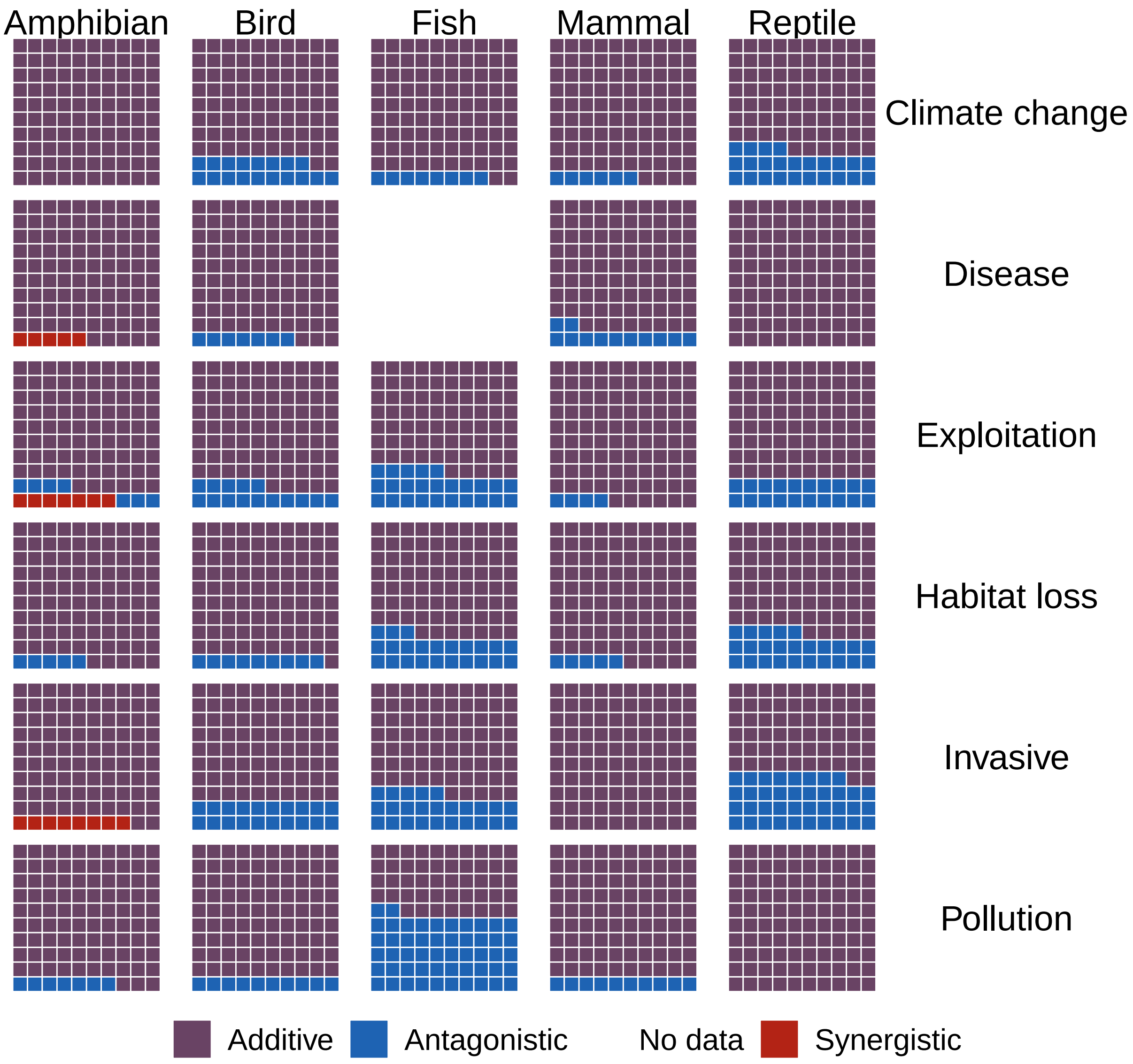


Figure S7: Spread of interactive effects across threats and taxa

Table S5: Proportion of threat interaction types across taxa. n represents the number of interaction types present. Frequency represents the proportion of that given interaction type for that threat and taxa

| Threat | Taxon | Interaction type | n | Frequency |
|---|---|---|---|---|
| Climate change | Amphibian | Synergistic | 0 | 0.000 |
| Climate change | Amphibian | Antagonistic | 0 | 0.000 |
| Climate change | Amphibian | Additive | 36 | 1.000 |
| Climate change | Bird | Synergistic | 0 | 0.000 |
| Climate change | Bird | Antagonistic | 15 | 0.179 |
| Climate change | Bird | Additive | 69 | 0.821 |
| Climate change | Fish | Synergistic | 0 | 0.000 |
| Climate change | Fish | Antagonistic | 3 | 0.083 |
| Climate change | Fish | Additive | 33 | 0.917 |
| Climate change | Mammal | Synergistic | 0 | 0.000 |
| Climate change | Mammal | Antagonistic | 2 | 0.056 |
| Climate change | Mammal | Additive | 34 | 0.944 |
| Climate change | Reptile | Synergistic | 0 | 0.000 |
| Climate change | Reptile | Antagonistic | 5 | 0.238 |
| Climate change | Reptile | Additive | 16 | 0.762 |
| Disease | Amphibian | Synergistic | 3 | 0.045 |
| Disease | Amphibian | Antagonistic | 0 | 0.000 |
| Disease | Amphibian | Additive | 63 | 0.955 |
| Disease | Bird | Synergistic | 0 | 0.000 |
| Disease | Bird | Antagonistic | 6 | 0.071 |
| Disease | Bird | Additive | 78 | 0.929 |
| Disease | Fish | Synergistic | NA | NA |
| Disease | Fish | Antagonistic | NA | NA |
| Disease | Fish | Additive | NA | NA |
| Disease | Mammal | Synergistic | 0 | 0.000 |
| Disease | Mammal | Antagonistic | 6 | 0.118 |
| Disease | Mammal | Additive | 45 | 0.882 |
| Disease | Reptile | Synergistic | 0 | 0.000 |
| Disease | Reptile | Antagonistic | 0 | 0.000 |
| Disease | Reptile | Additive | 21 | 1.000 |
| Exploitation | Amphibian | Synergistic | 3 | 0.071 |
| Exploitation | Amphibian | Antagonistic | 3 | 0.071 |
| Exploitation | Amphibian | Additive | 36 | 0.857 |
| Exploitation | Bird | Synergistic | 0 | 0.000 |
| Exploitation | Bird | Antagonistic | 15 | 0.147 |
| Exploitation | Bird | Additive | 87 | 0.853 |
| Exploitation | Fish | Synergistic | 0 | 0.000 |
| Exploitation | Fish | Antagonistic | 13 | 0.255 |
| Exploitation | Fish | Additive | 38 | 0.745 |
| Exploitation | Mammal | Synergistic | 0 | 0.000 |
| Exploitation | Mammal | Antagonistic | 3 | 0.040 |
| Exploitation | Mammal | Additive | 72 | 0.960 |
| Exploitation | Reptile | Synergistic | 0 | 0.000 |
| Exploitation | Reptile | Antagonistic | 13 | 0.197 |
| Exploitation | Reptile | Additive | 53 | 0.803 |
| Habitat loss | Amphibian | Synergistic | 0 | 0.000 |
| Habitat loss | Amphibian | Antagonistic | 3 | 0.053 |
| Habitat loss | Amphibian | Additive | 54 | 0.947 |
| Habitat loss | Bird | Synergistic | 0 | 0.000 |
| Habitat loss | Bird | Antagonistic | 11 | 0.092 |
| Habitat loss | Bird | Additive | 109 | 0.908 |
| Habitat loss | Fish | Synergistic | 0 | 0.000 |
| Habitat loss | Fish | Antagonistic | 14 | 0.233 |
| Habitat loss | Fish | Additive | 46 | 0.767 |
| Habitat loss | Mammal | Synergistic | 0 | 0.000 |
| Habitat loss | Mammal | Antagonistic | 5 | 0.054 |
| Habitat loss | Mammal | Additive | 88 | 0.946 |
| Habitat loss | Reptile | Synergistic | 0 | 0.000 |
| Habitat loss | Reptile | Antagonistic | 13 | 0.255 |
| Habitat loss | Reptile | Additive | 38 | 0.745 |

Table S5: Proportion of threat interaction types across taxa. n represents the number of interaction types present. Frequency represents the proportion of that given interaction type for that threat and taxa *(continued)*

| Threat | Taxon | Interaction type | n | Frequency |
|---|---|---|---|---|
| Invasive | Amphibian | Synergistic | 3 | 0.083 |
| Invasive | Amphibian | Antagonistic | 0 | 0.000 |
| Invasive | Amphibian | Additive | 33 | 0.917 |
| Invasive | Bird | Synergistic | 0 | 0.000 |
| Invasive | Bird | Antagonistic | 20 | 0.196 |
| Invasive | Bird | Additive | 82 | 0.804 |
| Invasive | Fish | Synergistic | 0 | 0.000 |
| Invasive | Fish | Antagonistic | 9 | 0.250 |
| Invasive | Fish | Additive | 27 | 0.750 |
| Invasive | Mammal | Synergistic | 0 | 0.000 |
| Invasive | Mammal | Antagonistic | 0 | 0.000 |
| Invasive | Mammal | Additive | 57 | 1.000 |
| Invasive | Reptile | Synergistic | 0 | 0.000 |
| Invasive | Reptile | Antagonistic | 8 | 0.381 |
| Invasive | Reptile | Additive | 13 | 0.619 |
| Pollution | Amphibian | Synergistic | 0 | 0.000 |
| Pollution | Amphibian | Antagonistic | 3 | 0.071 |
| Pollution | Amphibian | Additive | 39 | 0.929 |
| Pollution | Bird | Synergistic | 0 | 0.000 |
| Pollution | Bird | Antagonistic | 9 | 0.097 |
| Pollution | Bird | Additive | 84 | 0.903 |
| Pollution | Fish | Synergistic | 0 | 0.000 |
| Pollution | Fish | Antagonistic | 11 | 0.524 |
| Pollution | Fish | Additive | 10 | 0.476 |
| Pollution | Mammal | Synergistic | 0 | 0.000 |
| Pollution | Mammal | Antagonistic | 6 | 0.100 |
| Pollution | Mammal | Additive | 54 | 0.900 |
| Pollution | Reptile | Synergistic | 0 | 0.000 |
| Pollution | Reptile | Antagonistic | 0 | 0.000 |
| Pollution | Reptile | Additive | 36 | 1.000 |

Table S6: Proportion of threat interaction types across systems. n represents the number of interaction types present. Frequency represents the proportion of that given interaction type for that threat and system.

| Threat | System | Interaction type | n | Frequency |
|---|---|---|---|---|
| Climate change | Freshwater | Synergistic | 0 | 0.000 |
| Climate change | Freshwater | Antagonistic | 8 | 0.121 |
| Climate change | Freshwater | Additive | 58 | 0.879 |
| Climate change | Marine | Synergistic | 0 | 0.000 |
| Climate change | Marine | Antagonistic | 12 | 0.174 |
| Climate change | Marine | Additive | 57 | 0.826 |
| Climate change | Terrestrial | Synergistic | 0 | 0.000 |
| Climate change | Terrestrial | Antagonistic | 6 | 0.091 |
| Climate change | Terrestrial | Additive | 60 | 0.909 |
| Disease | Freshwater | Synergistic | 0 | 0.000 |
| Disease | Freshwater | Antagonistic | 13 | 0.188 |
| Disease | Freshwater | Additive | 56 | 0.812 |
| Disease | Marine | Synergistic | 0 | 0.000 |
| Disease | Marine | Antagonistic | 23 | 0.333 |
| Disease | Marine | Additive | 46 | 0.667 |
| Disease | Terrestrial | Synergistic | 10 | 0.133 |
| Disease | Terrestrial | Antagonistic | 3 | 0.040 |
| Disease | Terrestrial | Additive | 62 | 0.827 |
| Exploitation | Freshwater | Synergistic | 0 | 0.000 |
| Exploitation | Freshwater | Antagonistic | 7 | 0.093 |
| Exploitation | Freshwater | Additive | 68 | 0.907 |
| Exploitation | Marine | Synergistic | 0 | 0.000 |
| Exploitation | Marine | Antagonistic | 31 | 0.304 |
| Exploitation | Marine | Additive | 71 | 0.696 |
| Exploitation | Terrestrial | Synergistic | 9 | 0.107 |
| Exploitation | Terrestrial | Antagonistic | 0 | 0.000 |
| Exploitation | Terrestrial | Additive | 75 | 0.893 |
| Habitat loss | Freshwater | Synergistic | 0 | 0.000 |
| Habitat loss | Freshwater | Antagonistic | 21 | 0.206 |
| Habitat loss | Freshwater | Additive | 81 | 0.794 |
| Habitat loss | Marine | Synergistic | 0 | 0.000 |
| Habitat loss | Marine | Antagonistic | 20 | 0.196 |
| Habitat loss | Marine | Additive | 82 | 0.804 |
| Habitat loss | Terrestrial | Synergistic | 0 | 0.000 |
| Habitat loss | Terrestrial | Antagonistic | 6 | 0.050 |
| Habitat loss | Terrestrial | Additive | 114 | 0.950 |
| Invasive | Freshwater | Synergistic | 0 | 0.000 |
| Invasive | Freshwater | Antagonistic | 15 | 0.294 |
| Invasive | Freshwater | Additive | 36 | 0.706 |
| Invasive | Marine | Synergistic | 0 | 0.000 |
| Invasive | Marine | Antagonistic | 32 | 0.344 |
| Invasive | Marine | Additive | 61 | 0.656 |
| Invasive | Terrestrial | Synergistic | 9 | 0.107 |
| Invasive | Terrestrial | Antagonistic | 3 | 0.036 |
| Invasive | Terrestrial | Additive | 72 | 0.857 |
| Pollution | Freshwater | Synergistic | 0 | 0.000 |
| Pollution | Freshwater | Antagonistic | 14 | 0.187 |
| Pollution | Freshwater | Additive | 61 | 0.813 |
| Pollution | Marine | Synergistic | 0 | 0.000 |
| Pollution | Marine | Antagonistic | 24 | 0.286 |
| Pollution | Marine | Additive | 60 | 0.714 |
| Pollution | Terrestrial | Synergistic | 7 | 0.093 |
| Pollution | Terrestrial | Antagonistic | 0 | 0.000 |
| Pollution | Terrestrial | Additive | 68 | 0.907 |

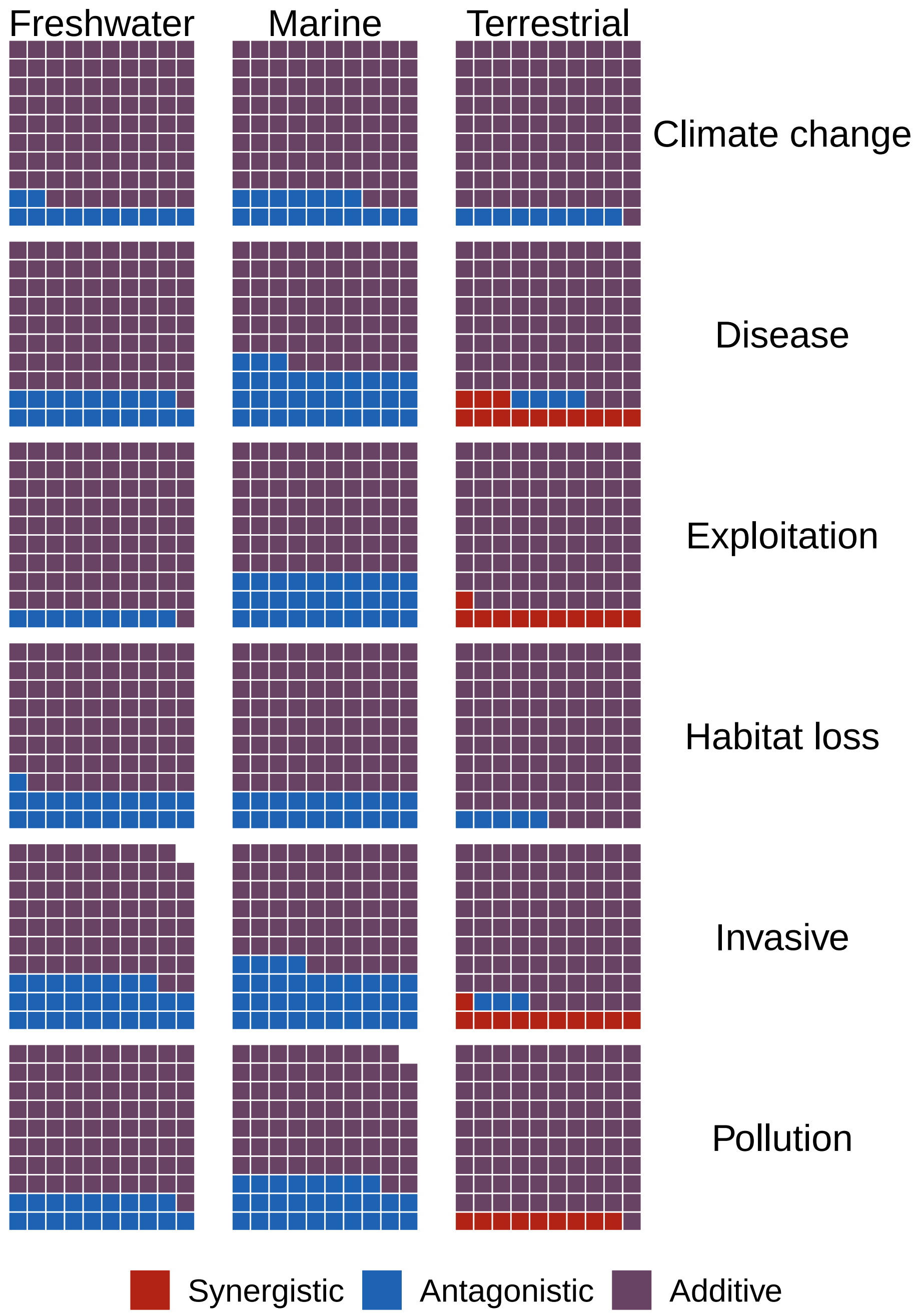


Figure S8: Spread of interactive effects across threats and systems.

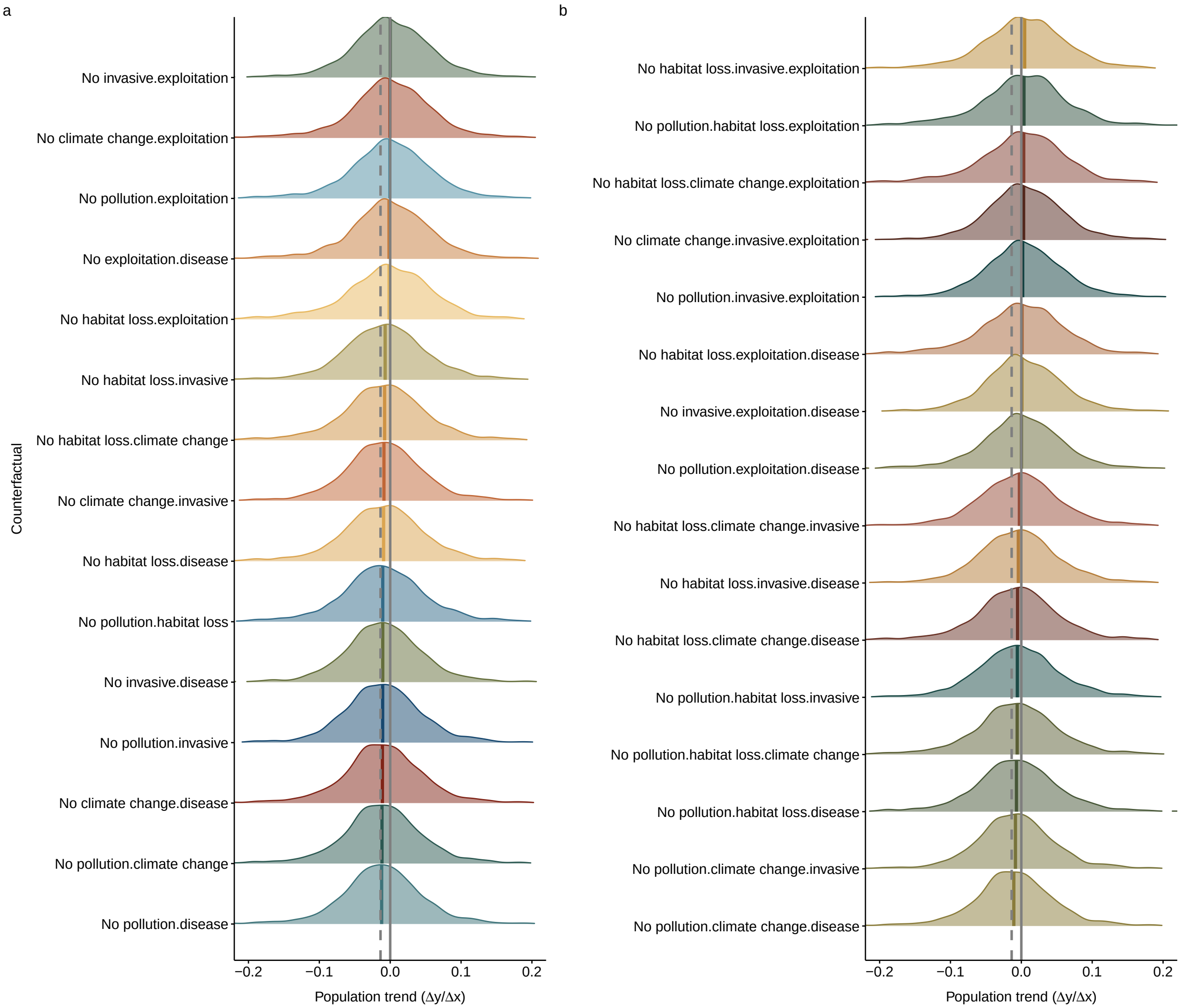


Figure S9: The counterfactual scenarios for multiple threats. The counterfactual scenarios represent the changes in the population trends of the 1,740 vertebrate time-series affected by threats, had there been different combinations of multiple threats. (a) Scenarios representing the global vertebrate population growth where different combinations of two threats were removed. (b) Scenarios representing the global vertebrate population growth where three threats were removed. The black line represents when the population trend is 0. The dotted line is the median trend of the threatened populations without any of the counterfactual scenarios.

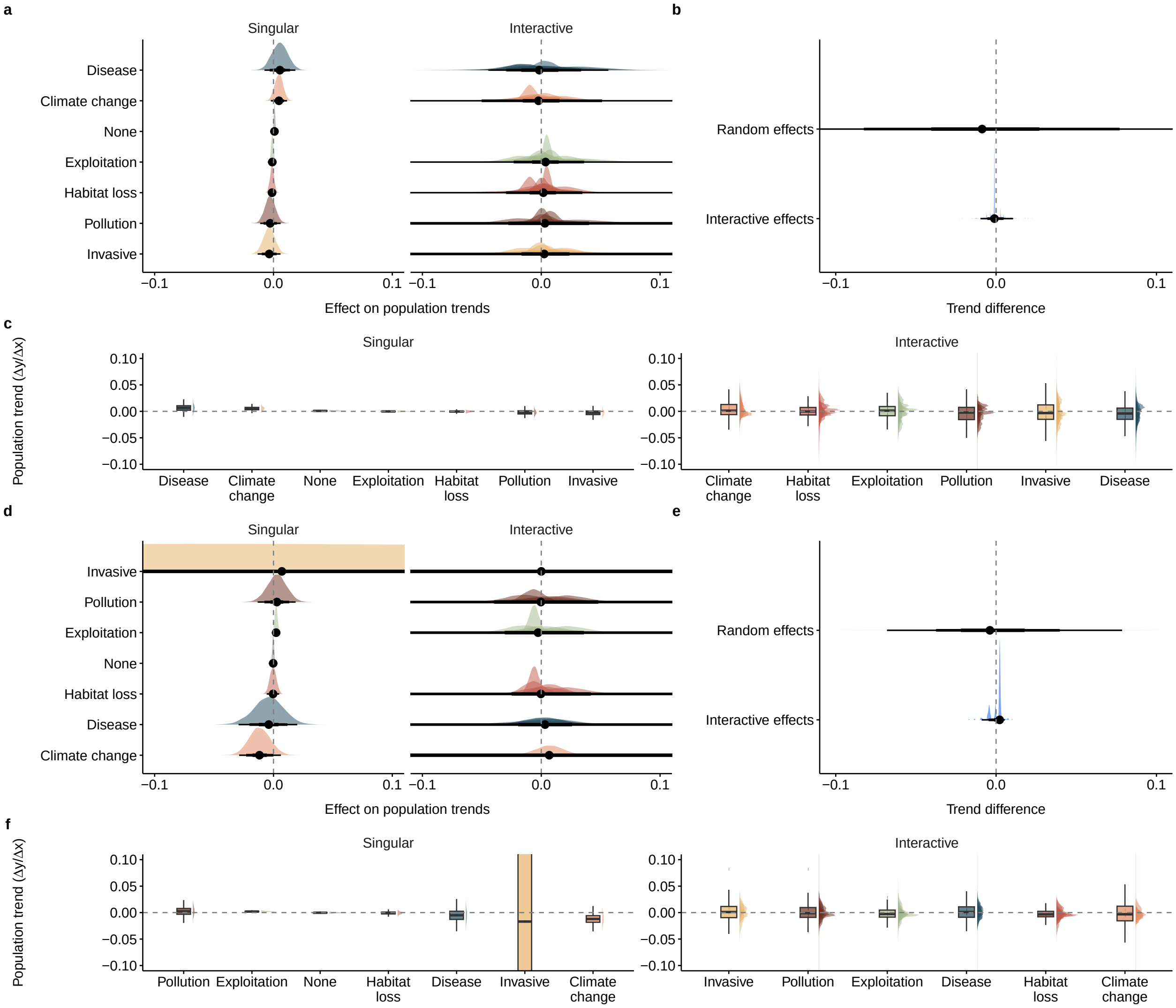


Figure S10: Influence of time series length on the relationship between threats and Living Planet Database trends. (a-c) Model estimates for time series containing 10 years of data. (d-f) Model estimates for time series containing 20 years of data.

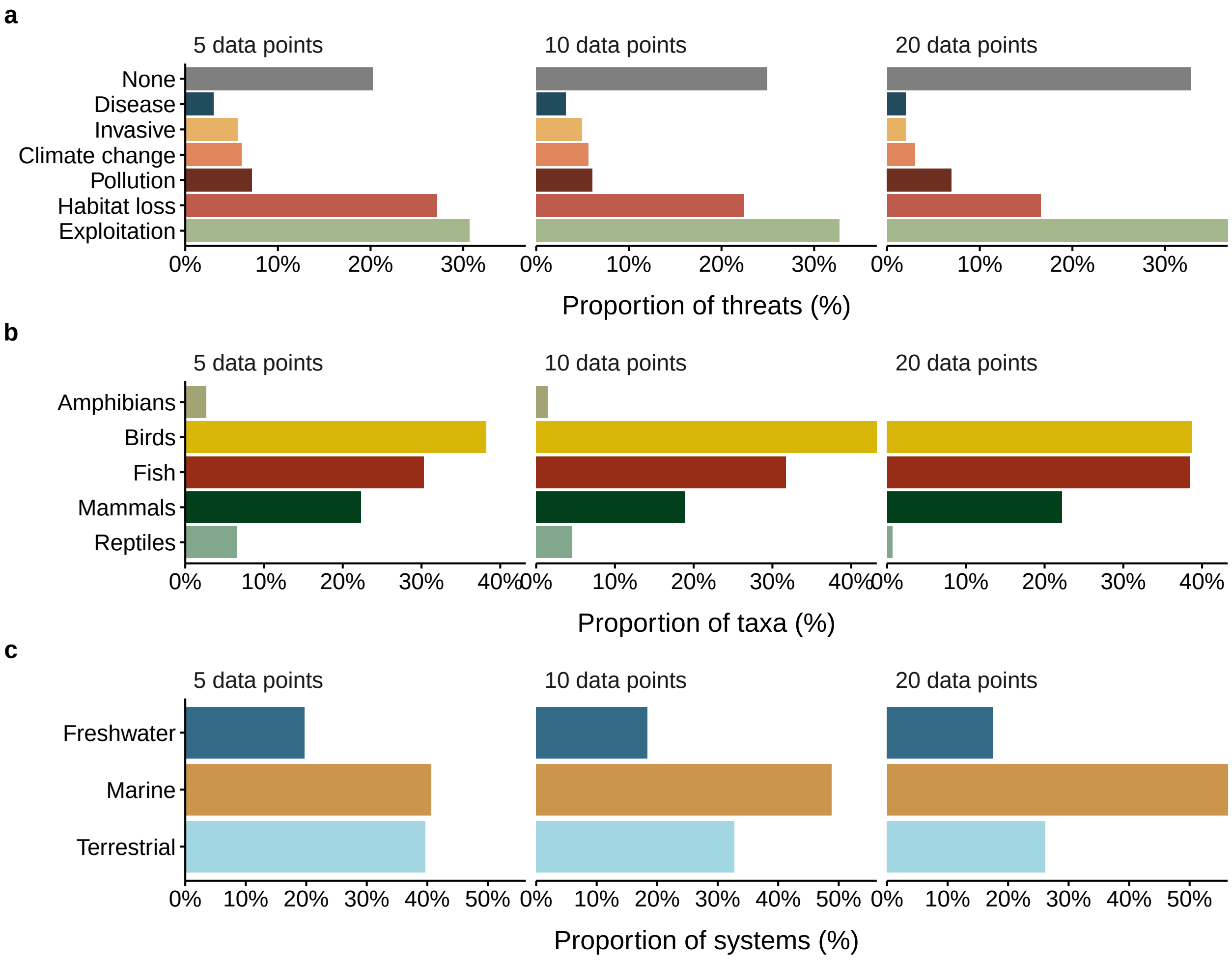


Figure S11: Frequency of threats, systems and taxa in the Living Planet Database once records have been filtered to time series containing 5, 10 and 20 years of data.

### 3 Appendix S3: Model checks

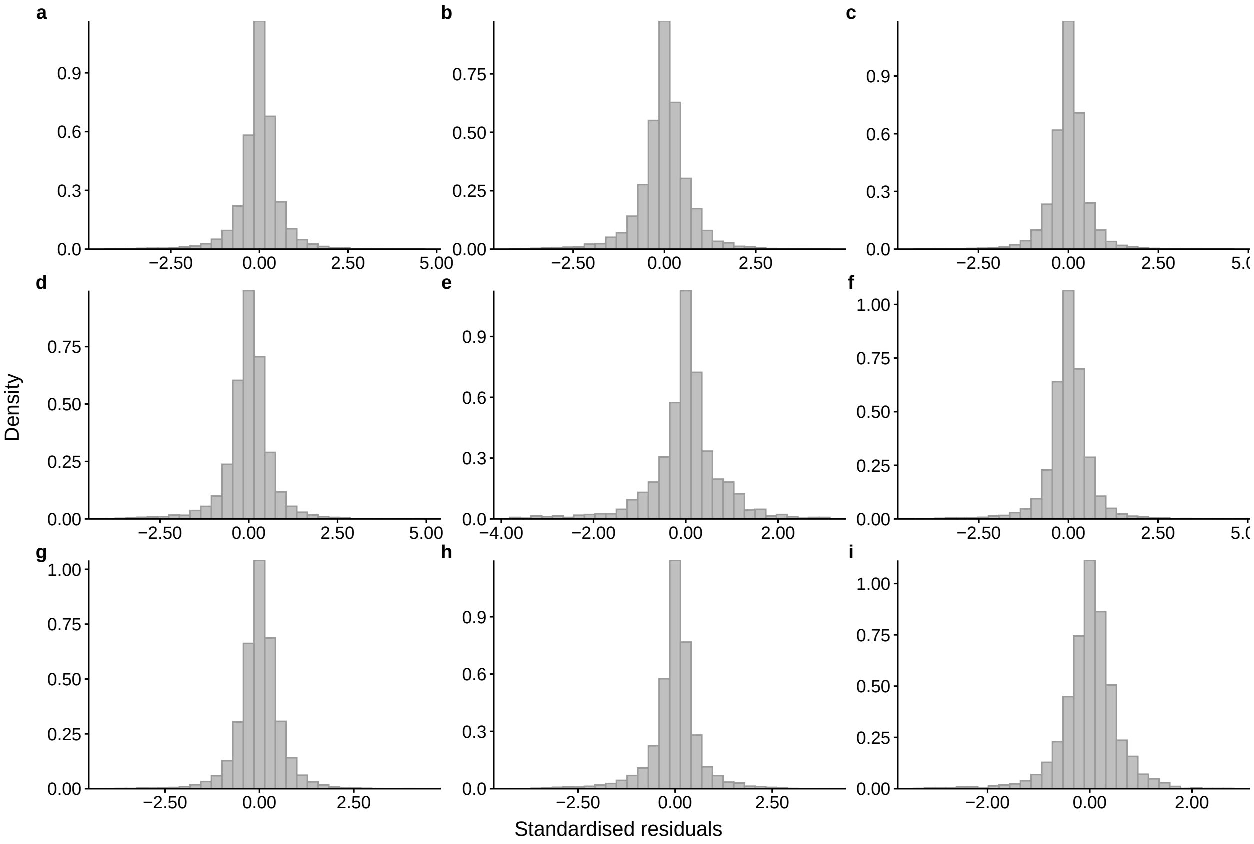


Figure S12: Distribution of the standardised residuals for the multilevel Bayesian models. Residuals were examined for the (a) general model, (b-d) system specific models (freshwater, marine, terrestrial), and (e-i) amphibians, birds, fishes, mammals, and reptiles respectively.

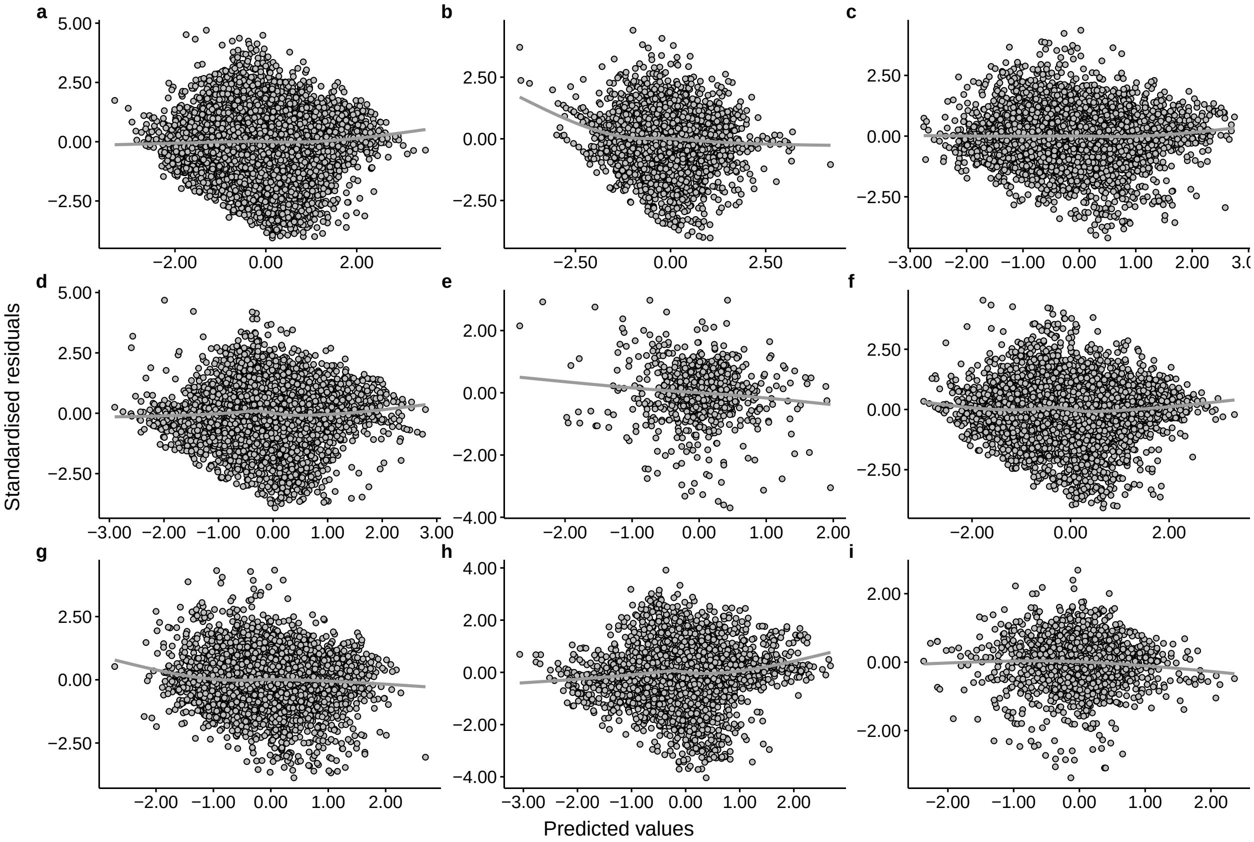


Figure S13: The standardised residuals vs the predicted values for the multilevel Bayesian models. The standardised residuals vs the predicted values have similar variances for all models, suggesting that the equal variance assumption is met. Residuals were examined for the (a) general model, (b-d) system specific models (freshwater, marine, terrestrial), and (e-i) amphibians, birds, fishes, mammals, and reptiles respectively.

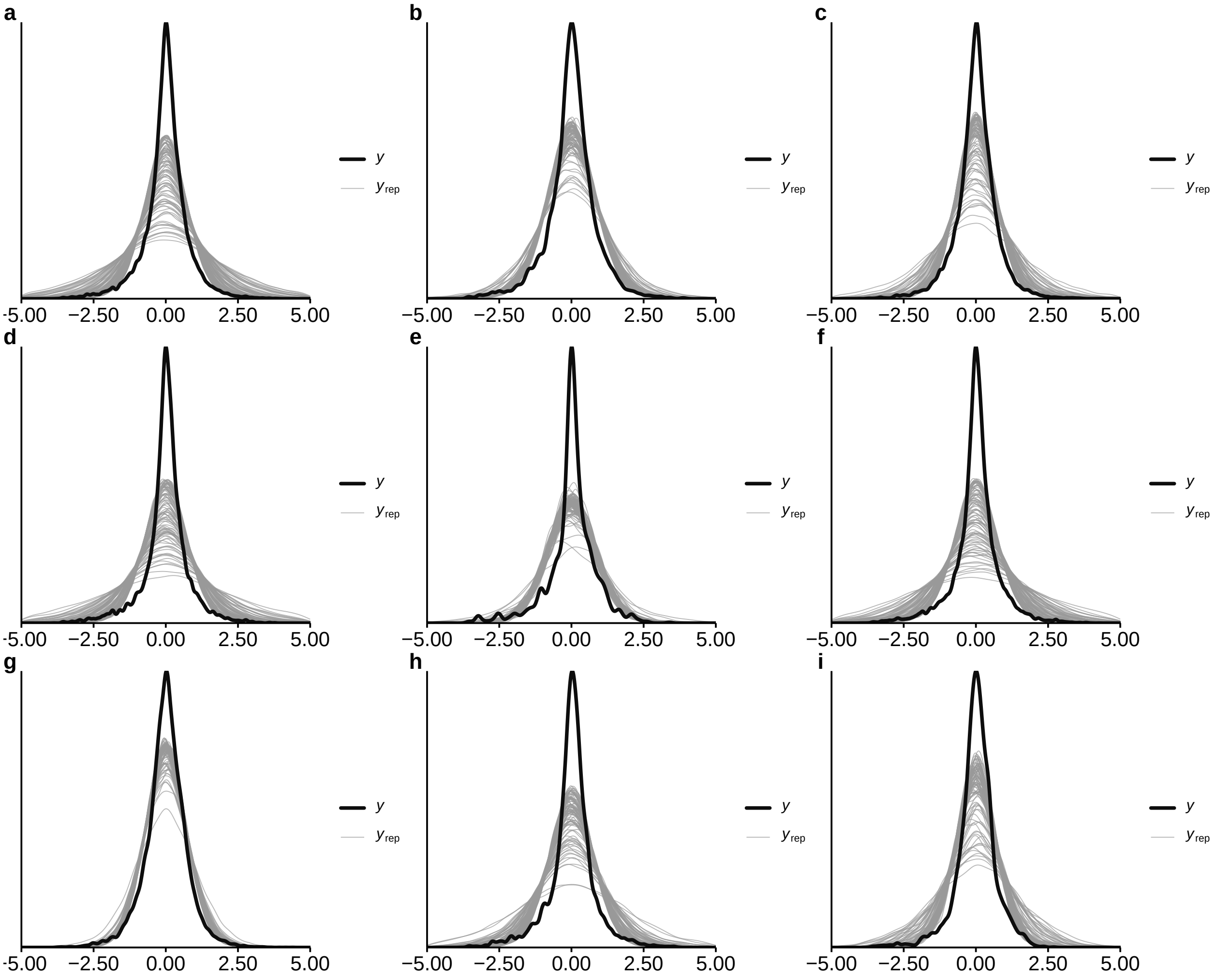


Figure S14: The posterior predictive checks for the multilevel Bayesian models. The posterior predictive checks do not show strong discrepancies between our data (dark lines, y) from the predictions from the model (light grey lines, yrep) for any of the models. However, the model shows a slight underestimation of the true zero values. The posterior predictive checks were examined for the (a) general model, (b-d) system specific models (freshwater, marine, terrestrial), and (e-i) amphibians, birds, fishes, mammals, and reptiles respectively.

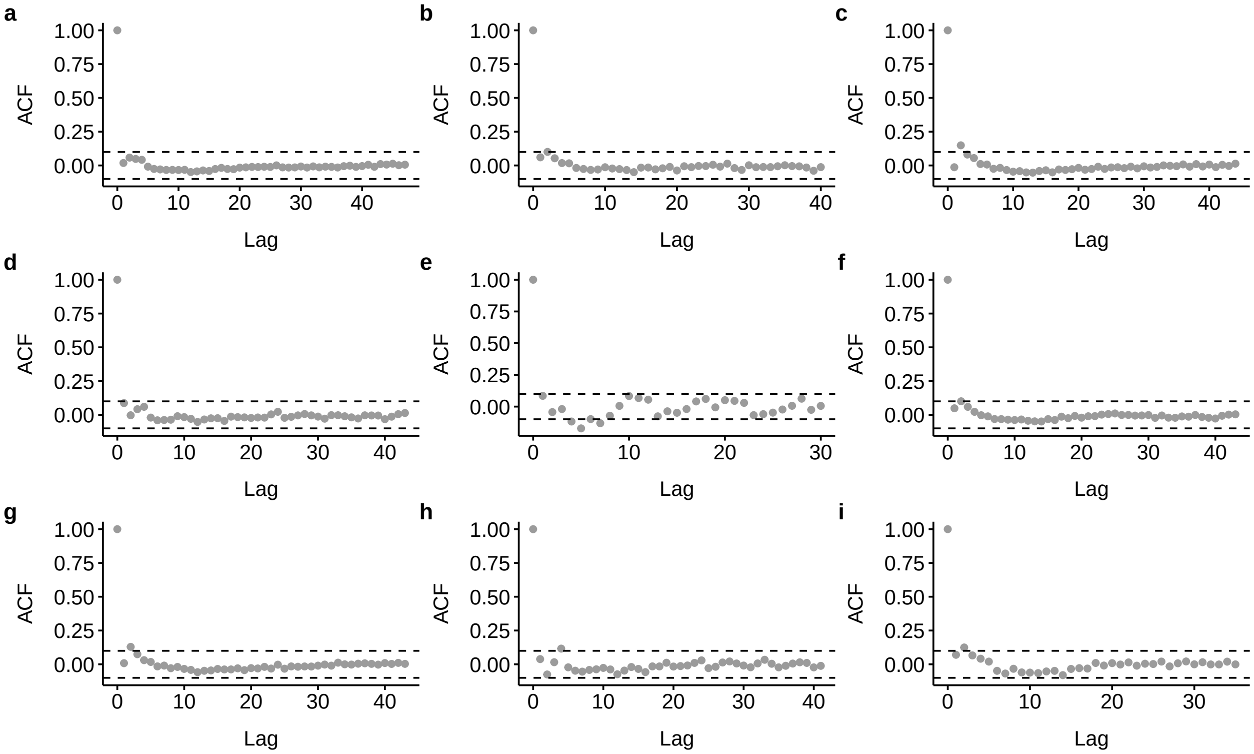


Figure S15: Autocorrelation of the standardised residuals for the multilevel Bayesian models. Limited evidence for autocorrelation is present for the (a) general model, (b-d) system specific models (freshwater, marine, terrestrial), and (e-i) amphibians, birds, fishes, mammals, and reptiles respectively.
